# AgRP Neuronal Suppression by Spontaneous Movement Encodes Exercise Reinforcement

**DOI:** 10.1101/2025.08.06.668910

**Authors:** Junichi Yoshida, Naofumi Suematsu, Shogo Soma, Ifat Levy, Tamas L. Horvath

## Abstract

Hypothalamic agouti-related peptide (AgRP) neurons that drive hunger-related behaviors are activated during energy deficit, encode negative valence but can be inhibited even by perception of food without digestion. Here, we show that in head-fixed mice, discrete motor actions such as spontaneous walking and conditioned licking rapidly suppress AgRP neuron activity. This suppression persisted throughout movement and terminated upon cessation, which was independent of food perception or conditioned cues. The degree of suppression was positively correlated with movement vigor. While optogenetic enhancement of AgRP neuron suppression did not initiate movement, it promoted reinforcement of concurrent walking behavior. These findings unmask that spontaneous, low intensity physical activity can reduce AgRP neuron activity reinforcing concurrent movement/exercise with implications for systemic health benefits.

## Main Text

Animals, including humans, engage in physical activity to acquire food when experiencing energy deficit. On the other hand, humans and animals can engage in excessive physical activity unrelated to food acquisition even though such activity consumes remaining calories and exacerbates hunger stress. In extreme cases, such as anorexia nervosa, patients compulsively exercise despite severe calorie restriction. How the brain resolves this paradoxical conflict and repeats the movements/exercise (sometimes compulsively) remains unknown.

Previous studies have shown that agouti-related peptide (AgRP)-expressing neurons in the hypothalamus expressing neuropeptide Y (NPY) and the inhibitory neurotransmitter, gamma amminobutytic acid (GABA) play a central role in hunger and satiety regulation and are essential for proper feeding behaviors^1–4^. These neurons are activated by calorie restriction^1,5–8^, which, in turn, triggers the release of stress hormones—such as corticosterone—via the hypothalamus-pituitary-adrenal (HPA) axis^9^. This suggests that calorie restriction is encoded as hunger stress through the activation of AgRP neurons. Upon activation, AgRP neurons not only promote feeding behaviors^10–15^ but also induce spontaneous, non-feeding-related movements, such as hyperlocomotion in confined spaces and compulsive grooming^16,17^. Typically, AgRP neurons are rapidly suppressed when animals perceive food (or food-predicting sensory cues) ^6,10,18,19^ and remain suppressed as nutritional signals enter the bloodstream during digestion^20^. Food perception inherently involves a sequence of movements, such as approaching food and engaging in licking or chewing. This raises the question of whether movement itself may contribute to AgRP neuronal suppression.

To examine the role of movement on the activity of AgRP neuronal suppression in negative energy balance (overnight food deprivation), we monitored AgRP neuronal activity during spontaneous wheel walking^21,22^ and conditioned licking responses^23,24^ using in vivo fiber photometry. We combined head-fixed behavioral paradigms with fiber photometry, which provides a high signal-to-noise ratio and enhances the detectability of movement-related effects on AgRP neurons.

### Spontaneous walking movement triggered the rapid suppression of AgRP neurons

We implemented a head-fixed behavioral paradigm (Fig. 1b & 2a) to precisely control walking and licking movements while simultaneously monitoring AgRP neuronal activity. Unlike free-moving conditions, head fixation minimizes extraneous behaviors such as rearing, grooming, and rotation, thereby reducing behavioral variability and ensuring a more controlled and reproducible experimental setting^21–24^. This approach also improved the signal-to-noise ratio in behavior-related fiber photometry signals. Z-scoring is commonly used to standardize recorded fiber photometry signals (see Methods for details). The accuracy of this calculation depends on the stability of the signal during the baseline period (e.g., several seconds before the behavior of interest), as it relies on the average and standard deviation of the signal within this period. If the signal fluctuates due to movements or if animals exhibit spontaneous behaviors during the baseline period, identifying movement-related fiber photometry signals becomes challenging. Under free-moving conditions, various behaviors (e.g., rearing and grooming) can occur during the baseline period, introducing variability. In contrast, head fixation reduces such extraneous movements, improving the signal-to-noise ratio in z-scored fiber photometry signals. This enhancement enabled us to reliably detect movement-driven AgRP neuronal activity.

**Figure 1.**
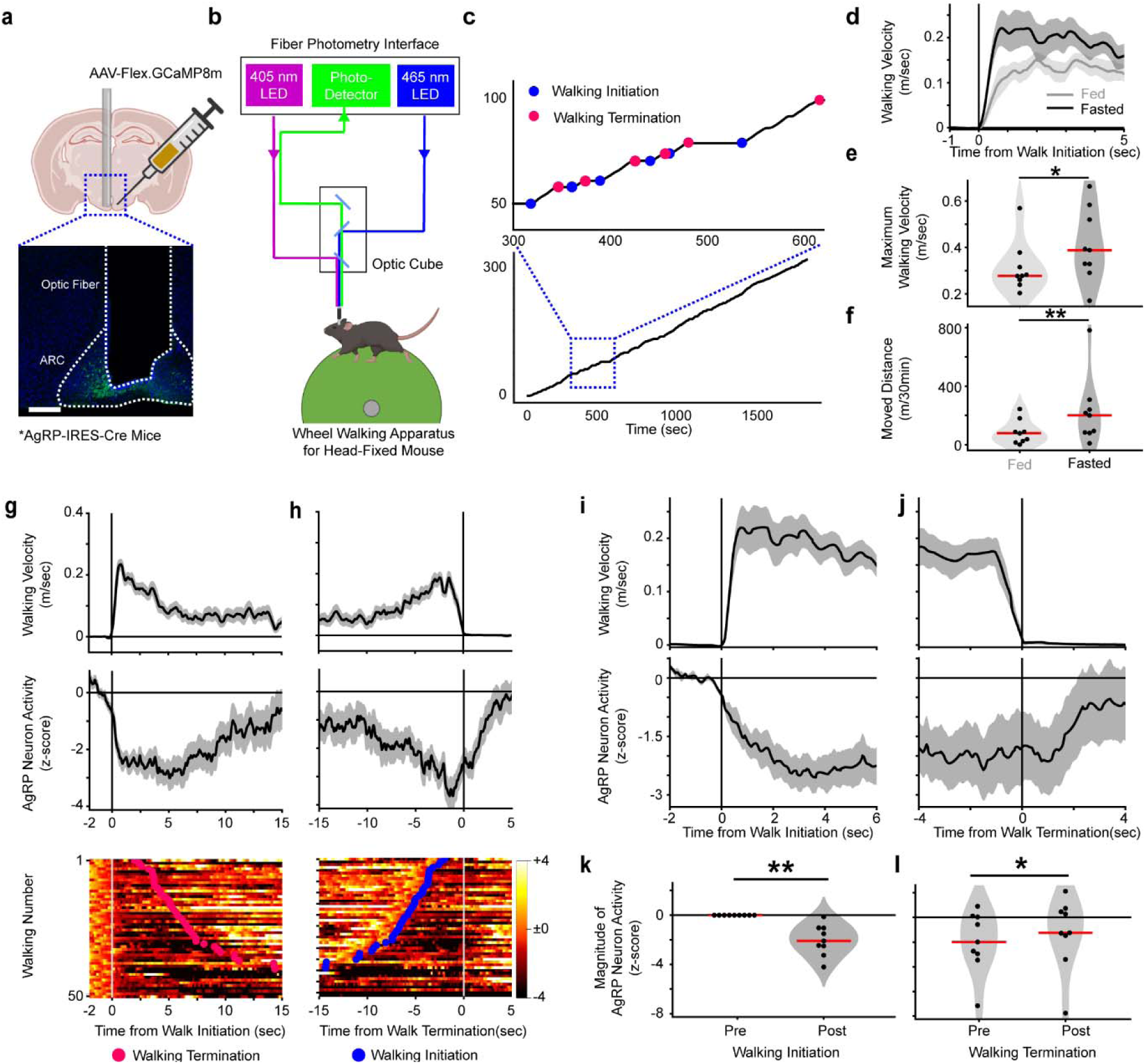
Spontaneous movements trigger rapid suppression in AgRP neurons. **(a**) Example of optic fiber placement and expression of the calcium indicators in AgRP neurons. Blue is DAPI and Green is GCaMP. The white bar indicates 200 µm. **(b)** Schematic of mice equipped for fiber photometry of AgRP neurons while walking on a wheel under head-fixed conditions. **(c)** Example of walking performance and identification of walking initiation/termination. Blue dots show walking initiation and red dots show walking termination. The top panel zooms in on the square in the bottom panel. **(d)** Average walking velocity dynamics of fed and fasted mice. **(e∼f)** Distribution of maximum walking velocity in the first 5 seconds (e) and moved distance in 30 minutes session (f). **(g∼h)** Example AgRP neuron activity at walking initiation (g) and termination (h). Top, average of walking velocity. Middle, average of z-scored AgRP neuron activity. Bottom, a trial-by-trial heat map of z-scored AgRP neuron activity. **(i∼j)** Average walking velocity (top), and AgRP neuron activity (bottom) among test animals at walking initiation (i) and termination (j). **(k∼l)** Distribution of AgRP neuronal suppression in pre-/post-window at walking initiation (k) and walking termination (l). In plots, data is presented as mean ± SEM. In plots and heat maps, bin time window is 50 ms. In violin plots, dots represent individual sessions and the red horizontal line indicates the median. ** means p < 0.01 for Wilcoxon-signed rank test against 0. N = 9 mice.

**Figure 2.**
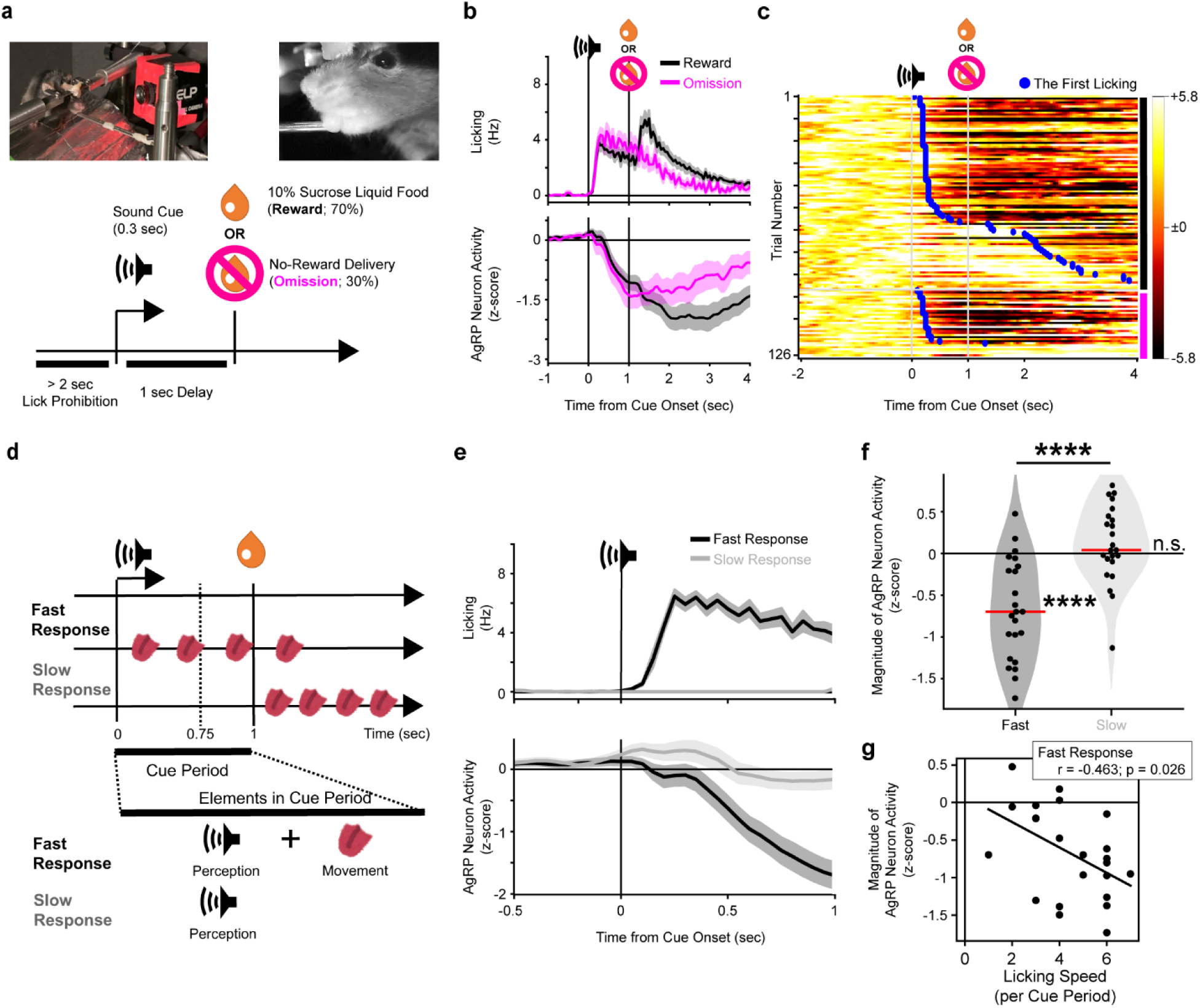
Rapid suppression in AgRP neuron is correlated to movement vigor. **(a**) Image of head-fixed Pavlovian conditioning task (top) and schematic design of the task (bottom). **(b)** Average licking (top) and AgRP neuron activity (bottom) among all sessions. **(c)** Heat map of AgRP neuron activity in an example session. The data is sorted along licking latency. Blue dots represent the first licking of each trial. **(d)** Schematic definition (left) and difference (right) of fast and slow response trials. **(e)** Average licking (top) and AgRP neuron activity (bottom) in fast and slow response trials. **(f)** Distribution of AgRP neuron activity magnitude during the cue period. **(g)** Correlation between licking number and AgRP neuron activity during the cue period in fast response trial. In plots, data is presented as mean ± SEM. In plots and heat maps, bin time window is 50 ms. In violin plots, dots represent individual session and the red horizontal line indicates the median. **** next of the red line means p < 0.0001 for Wilcoxon-signed rank test against 0. **** and ** between groups means p < 0.0001 and p < 0.01 for Wilcoxon-signed rank test to compare the trials. In the correlation plots, dots represent individual sessions and solid lines represent correlation line. The correlation lines, ρ, and p are estimated by Spearman rank-order correlation. N = 9 mice and n = 23 sessions.

In order to analyze AgRP neuron activity using fiber photometry, we targeted a calcium indicator (GCaMP8m^25^) to AgRP neurons and implanted an optic fiber unilaterally in the arcuate nucleus of the hypothalamus (Fig. 1a). Our goal was to measure AgRP neuron activity in mice spontaneously walking under conditions where they could not expect food acquisition. That’s why we used naive mice which had not yet been used for any experiments, such as Pavlovian task (Fig. 2-3). Mice were fasted overnight to activate AgRP neurons and were then placed on a wheel under head-fixed conditions with AgRP neuron fiber photometry for 30 minutes (Fig. 1b). In this setup, mice could walk spontaneously but only move forward or backward direction and were not able to rotate in any direction. We monitored the wheel’s rotation with a high-resolution rotary encoder (detecting 0.46 mm of movement minimally) connected to the wheel at a 4,000 Hz sampling rate, using the velocity of the wheel movement as a measure of the mice’s walking movement. Walking initiation was defined as occurring when the mice moved at more than 0.02 m/sec (or 1 mm per 50 ms) following a period of at least 3 seconds below this threshold. Walking termination was marked when the velocity dropped below the threshold for the subsequent 2 seconds (Fig. 1c). This head-fixed wheel apparatus allowed us to achieve more precise movement detection (1 mm per 50 ms) compared to standard free-moving tests (0.02 m per 100 ms under our experimental conditions; Extended Data Fig.1). This difference in movement detectability was critical for our study. Temporal and continuous suppressions in AgRP neurons were observed during food sensing and feeding under free-moving conditions (Extended Data Fig.1e, f, g, and h).

**Figure 3.**
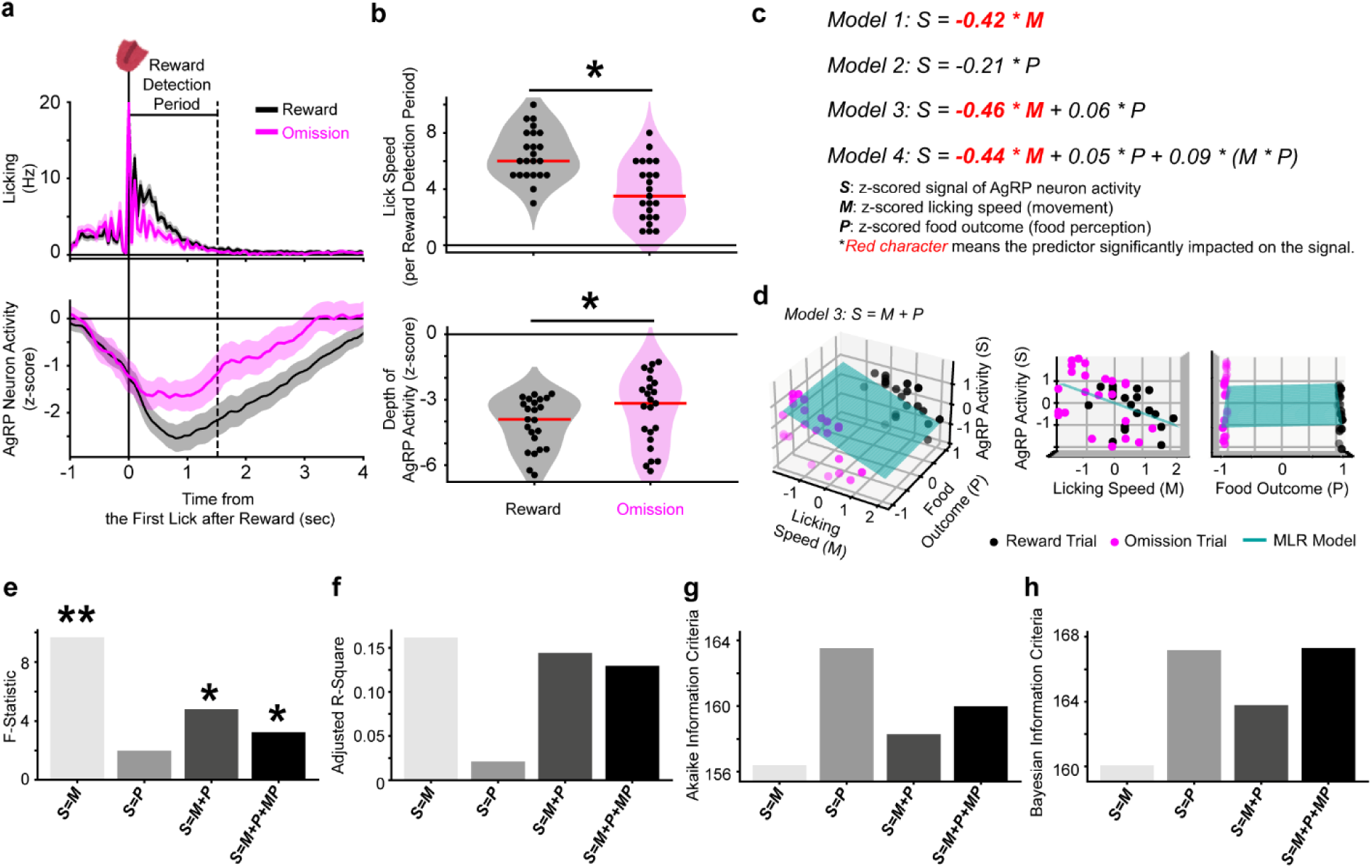
Movement is sufficient to explain rapid suppression in AgRP neurons at reward detection. **(a)** Average licking (top) and AgRP neuron activity (bottom) aligned with the first lick response after reward delivery (reward detection period). **(b)** Distribution of licking number (top) and AgRP neuron activity dip (bottom) during the reward detection period. **(c)** Four multiple linear regression (MLR) models estimated using AgRP neuron activity (S), licking number (M), food outcome (P), and the interaction between licking number and reward outcome (M*P). **(d)** 3D plots of MLR model 3 (S = M + P), with M-S and P-S planes shown in middle and right panels. **(e)** F-statistics, **(f)** adjusted R-squares, **(g)** Akaike information criteria, and **(h)** Bayesian information criteria of each MLR model. In panel **a**, data are presented as mean ± SEM with a 50 ms bin time window. In panel **b** violin plots, dots represent individual sessions, and the red horizontal line indicates the median. * denotes p < 0.05 in Wilcoxon-signed rank tests comparing reward and omission trials. MLR model coefficients in panel C were estimated via ordinary least squares, with significance tested using t-tests and p < 0.05 as the criterion. In panel D MLR plots, dots represent individual sessions, and the solid plane represents the estimated MLR model. * and ** in panel E denote p < 0.05 and p < 0.01 for F-tests. N = 9 mice, n = 23 sessions.

As previous studies have shown^26–28^, fasted mice exhibited more vigorous walking than fed mice, including in our head-fixed conditions (Fig. 1d; Fig. 1e; Maximum walking velocity in the first 5 seconds - Fed, Med = 0.18 [0.15, 0.20] m/sec; Fasted, Med = 0.26 [0.21, 0.30] m/sec; Wilcoxon signed-rank test, p = 0.008; Fig. 1f; Total moved distance in 30-minute session - Fed, Med = 78.70 [24.92, 88.45] m; Fasted, Med = 200.34 [84.00, 239.82] m; Wilcoxon signed-rank test, p = 0.008; N = 9 mice). Notably, this walking was independent of food availability. Fiber photometry revealed that AgRP neurons were significantly suppressed immediately after walking initiation (Movie1 and Fig. 1g, i, and k; pre-initiation, Med = 2.15*10^−17^[-5.02*10^−17^, 6.20*10^−17^] z-score; post-initiation, Med = -2.09 [- 2.50, -1.04] z-score; Wilcoxon signed-rank test, p = 0.004). This suppression of AgRP neurons was diminished after the end of walking (Fig. 1h, j, and l; pre-termination, Med = -1.99 [-2.95, 0.04] z-score; post-termination, Med = -1.24 [-1.47, 0.46] z-score; Wilcoxon signed-rank test, p = 0.020).

While the suppression magnitude in AgRP neuron activity was smaller during the walking than during food-feeding (Extended Data Fig.2), the walking-related suppression lasted during the entire walking period.

It may be that movement-driven AgRP neuronal suppression is a motion artifact. To address this, we monitored the Ca²^+^-independent signal using the isosbestic wavelength of the Ca²^+^ indicator and subtracted it from the Ca²^+^-dependent signal (Please see Methods for details). Although the Ca²^+^-independent signal exhibited some fluctuations, its magnitude was clearly lower than that of the Ca²^+^-dependent signal and did not show a suppression trend (Extended Data Fig.3). Additionally, we corrected all measured Ca²^+^-dependent signals using the Ca²^+^-independent signal and used the corrected signal as a measure of functional AgRP neuron activity (Extended Data Fig.4). To further interrogate this, we performed fiber photometry of proopiomelanocortin (POMC) neurons in the arcuate nucleus in head-fixed walking mice similarly as described above for AgRP neurons. In this case, in contrast to the response to AgRP neurons, we observed movement-driven activation rather than suppression of POMC neurons (Extended Data Fig.5). These observations are in line with the reported opposite activation pattern of POMC neurons than that of AgRP cells under various physiological conditions^18^ and the inhibitory tone of AgRP neurons on POMC cells^3^. Thus, we conclude that movement-driven AgRP neuronal suppression, as measured by fiber photometry, is not a motion artifact.

### Movement-driven AgRP neuronal suppression was correlated to movement vigor independent of food perception

To investigate whether the rapid suppression of AgRP neurons is triggered not only by walking but also by other movements, and whether this movement-driven suppression is influenced by food perception, we implemented a head-fixed Pavlovian conditioning task (Fig. 2a) following the head-fixed walking sessions. AgRP neuron activity was measured using fiber photometry, with a particular focus on its relationship to licking behavior. Licking responses were monitored using a lickometer based on resistance detection; however, this method only recorded licking when the mouse’s tongue made direct contact with the spout, potentially missing smaller licking movements. To overcome this limitation, we positioned a high-speed camera beside the mice’s faces to capture all licking events, taking advantage of the head-fixed setup to minimize extraneous movements. Using DeepLabCut^29^ for video analysis, we precisely tracked licking behavior and obtained a more accurate measure of licking vigor (Movie 1).

In the Pavlovian task (Fig. 2a), mice were fasted overnight before recording and placed on a locked wheel under head-fixed conditions. Each trial began when the mice inhibited licking for 2-3 seconds, with the exact duration randomly varied for each trial. The final 2 seconds of this licking prohibition period served as a baseline for z-scoring the fiber photometry signals. This rule improved the signal-to-noise ratio of z-scored AgRP neuron activity compared to sessions without it (data not shown). At the start of each trial, a reward-predictive cue (a 4 kHz pure tone for 0.3 seconds) was presented. A spout delivering a 10% sucrose solution as a liquid-food reward was positioned near the mouse’s mouth, with the reward delivered 1 second after the cue in 70% of the trials (reward trials). In the remaining 30% of trials, no reward was delivered after the cue (omission trials). Trials ended and transitioned to the next after a 3-second inter-trial interval if the mice licked the spout 1 second after the cue onset, ensuring no liquid-food residue remained on the spout when a new trial began.

Mice began licking immediately after the presentation of the conditioned cue, and a corresponding rapid suppression of AgRP neuron activity was observed (Fig. 2b and Movie 2). During the cue period, which spans from the onset of the conditioned cue to the reward delivery, both perceptual (from the conditioned cue) and motor (licking) components could potentially contribute to the suppression of AgRP neurons. To discern the primary factor driving this suppression, we visualized AgRP neuron activity on a trial-by-trial basis using a heat map and overlaid the timing of the first lick after the conditioned cue (Fig. 2c). This analysis revealed that the rapid suppression of AgRP neurons commenced following the initial licking movement, suggesting that the licking movement, rather than the perception of the conditioned cues, triggered the suppression. To interrogate this further, we analyzed AgRP neuron activity during the cue period by categorizing trials based on the latency of the licking response to the conditioned cue (Fig. 2d). Trials were divided into “fast response” trials, where the latency was less than 0.75 seconds, and “slow response” trials, where the latency was between 1 and 4 seconds. In the fast response trials, both the conditioned cue and licking occurred within the cue period, whereas in the slow response trials, only the cue was present, allowing for an evaluation of the relative contributions of licking and cue perception to AgRP neuronal suppression. In the fast response trials, AgRP neuron activity showed a significant decrease (Fig. 2e & f; fast trial, Med = - 0.69 [-1.12, -0.18] z-score; Wilcoxon signed-rank test, p = 2.6*10^−5^), while in the slow response trials, AgRP neuron activity did not exhibit a statistically significant change (Fig. 2e and f; slow trial, Med = 0.04 [-0.08, 0.42] z-score; Wilcoxon signed-rank test, p = 0.200). Furthermore, a comparison of neuronal activity magnitude between the two trial types indicated that AgRP neuron activity was significantly lower in fast response trials than in slow response trials (Fig. 2f, Wilcoxon signed-rank test, p = 1.0*10^−5^). In addition, AgRP neuron activity in fast response trials had a significant negative correlation to licking number (movement vigor) (Fig. 2g; Spearman rank-order correlation, ρ =-0.463, p = 0.026). These findings suggest that the licking movement is a major contributor to the suppression of AgRP neurons during the cue period.

The timing of the first lick after reward delivery corresponds to the perception of food taste (i.e., sweetness). To assess how food taste perception contributes to the rapid suppression of AgRP neurons, we compared the dynamics of AgRP neuron activity at the first lick after reward delivery between reward and omission trials (Fig. 3a). In reward trials, AgRP neuron activity showed significantly greater suppression than in omission trials (Fig. 3b bottom; reward trials, Med = -3.9 [-5.0, -3.1] z-score; omission trials, Med = -3.2 [-4.7, -2.3] z-score; Wilcoxon signed-rank test, p = 0.038). At the outset, this suggests that AgRP neuron activity is more suppressed by the perception of food.

However, mice exhibited more licking during this period in reward trials than in omission trials (Fig. 3b top; reward trials, Med = 6 [5, 8] licks; omission trials, Med = 3.5 [2, 5.5] licks; Wilcoxon signed-rank test, p = 3.3*10^−6^). Given that AgRP neuron activity was primarily suppressed by licking movement during the cue period (Fig. 2e and f), this raises the possibility that the greater suppression in AgRP neuron activity in reward trials might be due to more vigorous licking. To clarify the contributions of movement and perceptual elements to rapid AgRP neuron suppression at reward detection, we assessed the predictive power of each element using multiple linear regression (MLR) models. AgRP neuron activity, licking number, and food outcome (coded as 1 for reward and 0 for omission) were standardized by z-scoring and used to estimate four MLR models (Fig. 3c). An F-test revealed that models incorporating the movement parameter (licking number, M) showed at least one predictor significantly predicting AgRP neuron activity (S) (F-test; S=M, p = 0.003; S=M+P, p = 0.013; S=M+P+MP, p = 0.032), while the model using only the food perception parameter (food outcome, P) did not (Fig. 3e; F-test; S=P, p = 0.167). T-tests on each predictor confirmed that only movement (M) significantly contributed to AgRP neuronal suppression (Fig. 3c). The 3D plots of AgRP neuron activity, licking number, and food outcome confirmed a trend of the negative correlation between AgRP neuronal activity and licking number (M-S plane; Fig. 3d middle panel), but not with reward outcome (P-S plane; Fig. 3d right panel). This tendency of the negative correlation was further supported by Spearman correlations in both reward and omission trials (Extended Data Fig.6; reward trial, ρ =-0.361, p = 0.091; omission trial; ρ =-0.388, p = 0.067). The correlation estimated every session did not significantly differ in intercept (reward trial, Med = -3.2 [-4.3, -2.0]; omission trial, Med = -2.7 [-4.1, -1.24]; Wilcoxon signed-rank test, p = 0.134) or slope (reward, Med = -0.12 [-0.24, 0.01]; omission, Med = -0.08 [-0.31, 0.07]; Wilcoxon signed-rank test, p = 0.823) between conditions, indicating that movement-driven AgRP neuronal suppression was independent of food perception.

Finally, we compared model fit across the four MLR models (Fig. 3f–h) using adjusted R-squares, Akaike information criteria (AIC), and Bayesian information criteria (BIC). The model using only movement (M) scored best in all metrics, while the model using only food perception (P) scored worst. Adding food predictors (P and interaction MP) worsened model fit. These results indicate that AgRP neuronal suppression at reward detection window is primarily driven by movement vigor, independent of food perception.

Lastly, we assessed the termination of AgRP neuronal suppression. Analysis of AgRP neuron activity at the last lick following reward delivery indicated that suppression persisted until the final lick movement and began to return to baseline levels afterward (Extended Data Fig.7a). To statistically validate this observation, we calculated the rate of change in AgRP neuronal activity before and after the last lick (Extended Data Fig.7b and c). This analysis revealed that the rate of change in AgRP neuron activity was significantly negative until the end of licking (reward trial - pre-window, Med = - 1.05 [-1.54, -0.68] z-score/sec, Wilcoxon signed-rank test, p = 2.38*10^−7^; omission trial - pre-window, Med = -1.06 [-1.50, -0.29] z-score/sec, Wilcoxon signed-rank test, p = 2.10*10^−5^). Conversely, the rate of change in activity became significantly positive after the last lick (reward trial - post-window, Med = 0.94 [0.56, 1.32] z-score/sec, Wilcoxon signed-rank test, p = 2.38*10^−7^; omission trial - post-window, Med = 0.21 [0.03, 1.06] z-score/sec, Wilcoxon signed-rank test, p = 0.0017). The rate of change in AgRP neuron activity was significantly different before and after the last lick in both reward and omission trials (reward trial, p = 2.38*10^−7^; omission trials, Wilcoxon signed-rank test, p = 0.0001), supporting our finding from the wheel walking experiment (Fig. 1) that AgRP neuronal suppression started at the beginning of a movement and it terminated at the end of the movement.

### Amplification of the AgRP neuronal suppression promotes reinforcement of concurrent movement

Our study revealed that movement—even when unrelated to food acquisition or intake—rapidly suppresses AgRP neuron activity (Fig. 1–3). However, the magnitude of movement-driven suppression was significantly smaller than that induced by feeding (Extended Data Fig.1g, h). Specifically, movement caused approximately 30% of the suppression observed during feeding (Extended Data Fig.1h). What role does this smaller suppression play in behavior? Does it contribute to the execution or regulation of movement itself? Or does it alter the quality or persistence of movement in a meaningful way? To address these questions, we employed optogenetic manipulation to artificially induce or amplify AgRP neuronal suppression (Fig. 4–6).

**Figure 4.**
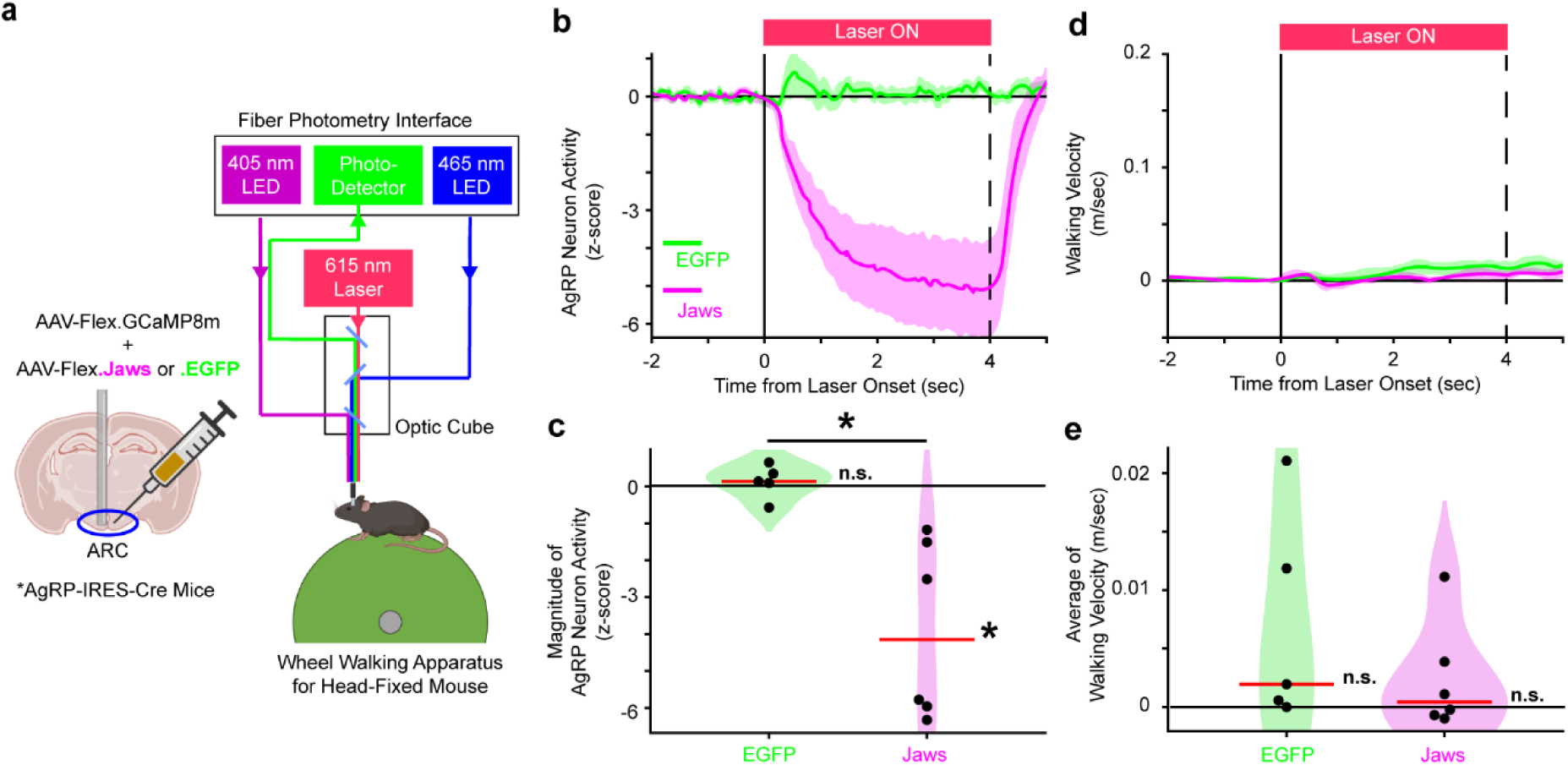
Optogenetic AgRP suppression does not evoke walking movement. (a) Schematic of virus injection and optic fiber implant at the ARC (left) and mice equipped for single fiber optogenetics-fiber-photometry of AgRP neurons while walking on a wheel under head-fixed conditions (right). **(b)** Average of AgRP neuron activity aligned at laser onset. **(c)** Distribution of AgRP neuron activity during the laser irradiation. **(d)** Average of walking velocity aligned at the laser onset. **(e)** Distribution of averaged walking velocity during the laser irradiation. In panel b, c, d, and e, green represents AgRP-EGFP mice and magenta represents AgRP-Jaws mice. In panel **b** and **d**, data are presented as mean ± SEM with a 50 ms bin time window. In violin plots, dots represent individual sessions and the red horizontal line indicates the median. * next of the red line means p < 0.05 for Wilcoxon-signed rank test against baseline 0. * between trials means p < 0.05 for Wilcoxon-signed rank test to compare the mouse group. N = 5 AgRP-EGFP mice and 6 AgRP-Jaws mice.

In the first experiment, we asked whether AgRP neuronal suppression is sufficient to initiate movement. We expressed either the inhibitory opsin Jaws or a control fluorescent protein (EGFP) in AgRP neurons (AgRP-Jaws or AgRP-EGFP mice). To make AgRP neurons be activated, calorie restricted mice were used as same as the fiber photometry experiments. Optogenetic stimulation was delivered during stationary periods on a running wheel (Fig. 4a), and AgRP neuron activity was simultaneously monitored using fiber photometry. We confirmed that laser stimulation effectively suppressed AgRP neuron activity in AgRP-Jaws mice (Fig. 4b, c; AgRP-EGFP, Med = 0.13 [0.09, 0.34] z-score/sec, Wilcoxon signed-rank test against baseline, p = 0.44; AgRP-Jaws, Med = -4.15 [-5.91, -1.76] z-score/sec, Wilcoxon signed-rank test against baseline, p = 0.03; Wilcoxon signed rank test for group comparison, p = 4.33*10^−4^). However, this suppression did not induce walking in either AgRP-Jaws or AgRP-EGFP mice (Fig. 4d, e; AgRP-EGFP, Med = 1.9*10^−4^ [5.6*10^−5^, 1.2*10^−3^] m/sec, Wilcoxon signed-rank test against baseline, p = 0.13; AgRP-Jaws, Med = 4.2*10^−5^ [-5.9*10^−5^, 3.2*10^−4^] m/sec, Wilcoxon signed-rank test against baseline, p = 0.44; Wilcoxon signed rank test for group comparison, p = 0.25), suggesting that suppression of AgRP neurons alone is not sufficient to initiate movement.

Next, we tested whether amplifying the natural, movement-driven suppression of AgRP neurons could alter ongoing movement (Fig. 5a). Using a closed-loop system, we delivered optogenetic stimulation only during wheel running (Fig. 5b). This design ensured that AgRP suppression was amplified exclusively during active movement, and only in AgRP-Jaws mice. On the first day of stimulation, there was no significant difference in walking speed between AgRP-Jaws and control mice (Fig. 5c, d; AgRP-EGFP, Med = 0.09 [0.03, 0.12] m/sec; AgRP-Jaws, Med = 0.07 [0.03, 0.14] m/sec; Wilcoxon signed rank test for group comparison, p = 0.63). However, by day 3, AgRP-Jaws mice walked significantly faster than controls (Fig. 5e, f; AgRP-EGFP, Med = 0.06 [0.02, 0.13] m/sec; AgRP-Jaws, Med = 0.17 [0.12, 0.32] m/sec; Wilcoxon signed rank test for group comparison, p = 0.04), indicating that movement vigor was reinforced by the amplified suppression of AgRP neurons. Consistent with this, AgRP-Jaws mice also showed a greater total distance traveled across the three-day period (Fig. 5g∼j; KS-test for group comparison – total accumulated moved distance through day 1 to day 3, p = 0.0054; day 1, p = 3.4*10^−3^; day 2, p = 0.07; day 3, p = 1.2*10^−3^) although original walking performances between AgRP-Jaws and AgRP-EGFP mice were not significantly different (Extended Data Fig.8). These findings support the notion that suppression of AgRP neuronal activity during movement reduces hunger-related stress and reinforces the accompanying behavior.

**Figure 5.**
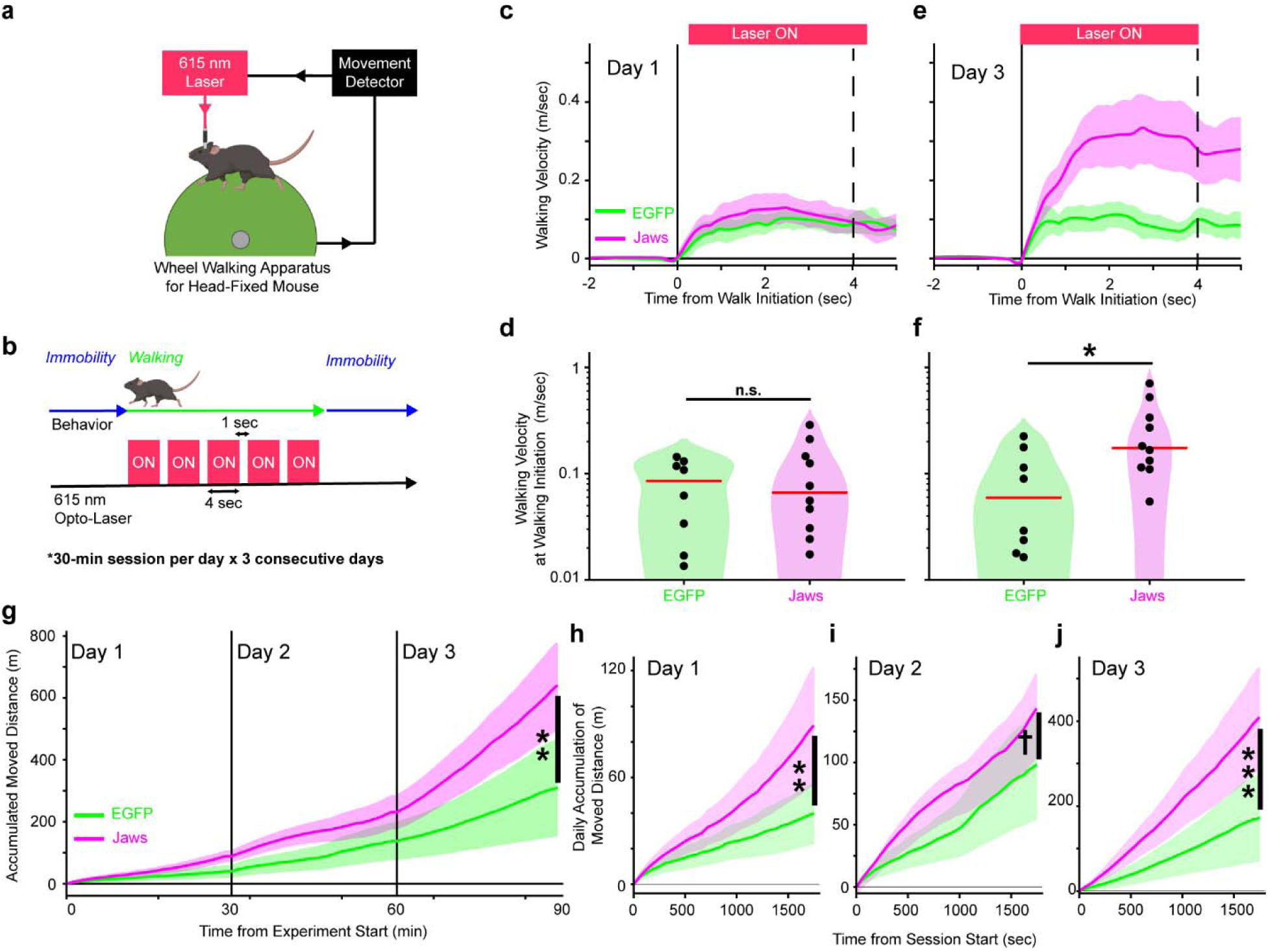
Optogenetic amplification of AgRP suppression promotes reinforcement of concurrent walking movement. **(a)** Schematic of the closed-loop system linking the opto-laser to the wheel. **(b)** Schematic of the opto-laser stimulation protocol. **(c, d)** Average **(c)** and distribution **(d)** of walking velocity on day 1. **(e, f)** Average **(e)** and distribution **(f)** of walking velocity on day 3. **(g)** Accumulated walking distance across three consecutive daily sessions. (**h–j**) Accumulated walking distance on day 1 **(h)**, day 2 **(i)**, and day 3 **(j)**. In panels **c–j**, green indicates AgRP-EGFP mice and magenta indicates AgRP-Jaws mice. In panels **c** and **e**, data are shown as mean ± SEM using a 50 ms time bin. In panels **g–j**, data are shown as mean ± SEM using a 50 s time bin. In violin plots, dots represent individual sessions and red horizontal lines indicate the median. ✝p < 0.1, *p < 0.05, and **p < 0.01 between groups by Wilcoxon signed-rank test (**d** and **f**) or Kolmogorov–Smirnov test (**h-j**). Except for a panel **g**, FDR Benjamini-Hochberg method was used for multiple comparison correction (**d, f, and h-j**). N = 8 AgRP-EGFP mice and 10 AgRP-Jaws mice.

**Figure 6.**
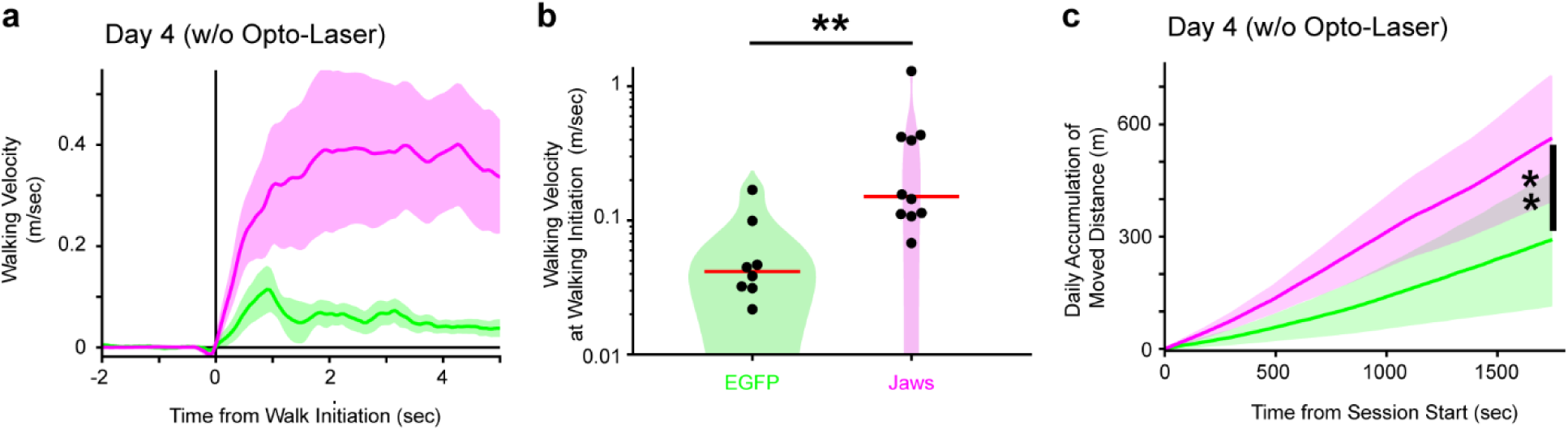
Walking reinforcement persists after cessation of optogenetic amplification of walking-driven AgRP suppression. **(a)** Average and **(b)** distribution of walking velocity on day 4, when no opto-laser stimulation was applied. **(c)** Average accumulated walking distance on day 4. Green indicates AgRP-EGFP mice and magenta indicates AgRP-Jaws mice. In panel **a**, data are shown as mean ± SEM using a 50 ms time bin. In panel **c**, data are shown as mean ± SEM using a 50 s time bin. In the violin plot **(b)**, dots represent individual mice and red horizontal lines indicate the median. **p < 0.01 between groups by Wilcoxon signed-rank test **(b)** or Kolmogorov–Smirnov test **(c) followed by** FDR Benjamini-Hochberg correction against day 1 ∼ 4 consecutive repeats of the statistical tests. N = 8 AgRP-EGFP mice and 10 AgRP-Jaws mice.

Importantly, this reinforcement persisted even after the optogenetic stimulation ended: AgRP-Jaws mice continued to exhibit increased walking velocity (Fig. 6a, b; AgRP-EGFP, Med = 0.04 [0.03, 0.06] m/sec; AgRP-Jaws, Med = 0.15 [0.11, 0.42] m/sec; Wilcoxon signed rank test for group comparison, p = 8.2*10^−3^) and greater cumulative movement (Fig. 6c; ; KS-test for group comparison, p = 3.4*10^−3^) compared to controls on day 4, when the laser for optogenetics was not triggered by walking performance at all.

Taken together, our data show that movement-driven suppression of AgRP neurons not only modulates immediate motor output but also facilitates long-term reinforcement of physical activity coinciding with hunger signal attenuation.

## Discussion

Here, we revealed how spontaneous, low intensity movements contribute to the rapid suppression of AgRP neurons, which persisted until movement termination. Under the Pavlovian conditioning task paradigm, AgRP neuronal suppression was primarily triggered by movement independent of food perception. The suppression of AgRP neurons correlated with movement vigor. Finally, amplification of the AgRP neuronal suppression synchronized with movement encoded a signal to promote movement reinforcement.

Earlier studies failed to detect movement-driven suppression in AgRP neurons in freely moving mice^19^. Our findings arose from head-fixed behavioral paradigm. As previously discussed, head fixation minimizes extraneous behaviors and improves the signal-to-noise ratio in behavior-related fiber photometry signals. This advantage is particularly important for detecting neuronal dynamics in populations encoding various movements, such as motor cortical pyramidal neurons^30,31^, striatal medium spiny neurons^32^, cerebellar Purkinje cells^33^, and AgRP neurons. In fact, we also failed to detect AgRP neuronal dynamics at movement initiation when using a free-moving treadmill^17^, where mice could engage in extraneous behaviors during the waiting stage, such as rearing, grooming, and sniffing—behaviors that may trigger AgRP neuronal suppression before the mouse begins running on the treadmill. While it is possible to observe movement-driven AgRP neuronal suppression even with a lower signal-to-noise ratio, the magnitude of suppression is smaller than that driven by other factors, such as food perception and nutrient signals (Extended Data Fig.2). This suggests that detecting these relatively small fluctuations in AgRP neuron activity during movement would be challenging without sufficient signal resolution.

Our findings highlight movement as a key factor driving AgRP neuronal suppression independent of food perception. While we demonstrated that motion artifacts do not account for this suppression, a related question arises: how does movement-driven suppression interact with other known modulators of AgRP activity, such as taste? One particularly complex case is the transient suppression of AgRP neurons during licking behavior, as licking involves both the motor action of licking and the sensory experience of tasting. Knight and colleagues recently reported that licking an empty spout did not evoke AgRP suppression, whereas licking a spout containing a sweet- or fat-tasting liquid did, leading them to conclude that taste, rather than movement, was the primary driver of suppression^19^. However, we propose that this difference in interpretation may stem from variations in the time scale of licking behavior. In their study, mice exhibited approximately 100 licks over 8–10 seconds, whereas in our experiment, mice performed fewer than 10 licks within 2 seconds (Fig. 2b). This suggests that our study captured the initiation of licking, while their study focused on its prolonged execution. Notably, Knight et al. also observed a small suppression during the first 5 seconds of empty-bottle licking—a time window that aligns with the duration of licking in our Pavlovian conditioning paradigm (Fig. 2b). Considering these points, we propose that micro-movements, such as licking, can evoke rapid AgRP suppression in a manner dependent on movement vigor, though this effect is short-lived compared to the suppression induced by larger-scale movements like walking (which occurs over 10–100 seconds). Furthermore, when food is present, taste likely contributes to AgRP suppression, particularly when licking is sustained for longer durations, such as more than 5 seconds.

The fact that movement itself triggers a rapid suppression of AgRP neurons and amplification of the AgRP neuronal suppression time-locked with a movement reinforces the concurrent movement suggests that movement can alleviate negative valance or stress associated with hunger, which works as a reinforcer for the movement (Fig. 7). When animals are hungry, they engage in behaviors like exploration to obtain food and alleviate their hunger. However, the accumulation of movement during these behaviors consumes calories in the body that could lead to further activation of AgRP neurons and an increase in negative-valence. In turn, movements that animals engage in to relieve hunger stress could exacerbate the stress. Instead, animals can initiate and sustain food-seeking behavior when hungry. Humans and animals can compulsively emgage in physical activity unrelated to food-seeking. Our findings may provide a mechanism via which this paradox can be resolved and subjects gain ability to continue physical activity under calorie restriction: movement partially suppresses the activity of AgRP neurons, thereby alleviating hunger-associated distress and also working as a reinforcer to the concurrent movements even though this suppression is smaller compared to when food is actually obtained (Extended Data Fig.2). Since this effect continues during movement (Fig. 1g to l, 2e to f and Extended Data Fig.7), animals are likely able to continue behaviors such as exploration until they find food. Additionally, the relationship between AgRP neurons and movement may explain why caloric restriction or experimental activation of AgRP neurons induces not only increased food intake but also various spontaneous movements (e.g., more locomotion and excessive grooming)^16^. Animals with activated AgRP neurons and increased hunger may repeatedly perform movements in their environment to partially suppress AgRP neurons and alleviate their negative valence during hunger.

**Figure 7.**
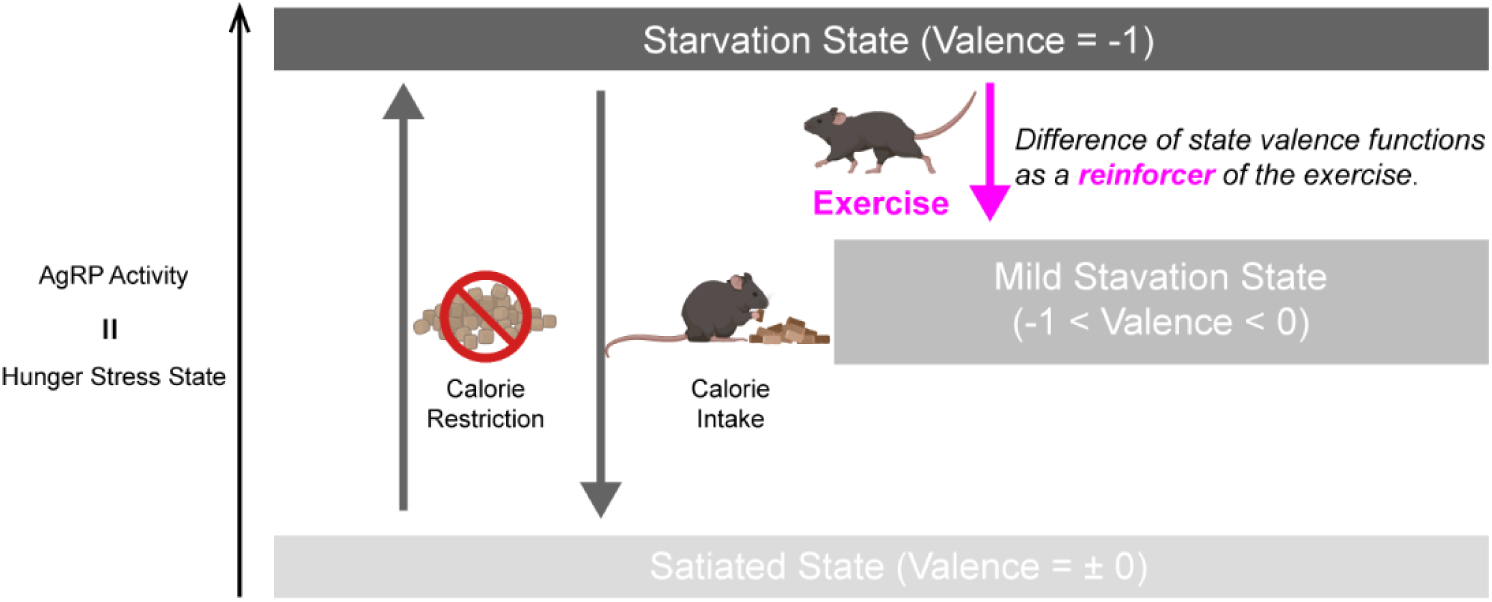
Schematic model of the role of AgRP neuron suppression in exercise reinforcement. This diagram illustrates a proposed mechanism by which physical activity under calorie restriction suppresses AgRP neuronal activity. The suppression attenuates the negative valence associated with hunger signals, reinforcing the ongoing movement. Repeated coupling of movement with AgRP suppression may contribute to the persistence and reinforcement of exercise behavior, even under energy-deficient conditions.

Our findings may also provide new insights into understanding of symptoms observed in subjects with anorexia nervosa (AN). Large number of subjects with AN are known to engage in hyperactivity/compulsive-exercise, even when they do not consume the necessary calories due to extreme dietary restrictions^34–41^. Some studies using rodent anorexia model (activity-based anorexia: ABA) reported that exercise during ABA paradigm would become positively reinforcing, potentially reducing the negative stress of hunger^27,42^. Our research revealed that movements suppress activity in AgRP neurons, which work as a reinforcer for the accompanied movements. Considering that many studies suggested the relationship between the AN and abnormal AgRP function (allelic variation in *agrp* genes^43,44^, AGRP peptide and mRNA level in the plasma^45–49^, and AgRP neuron activity^50^ in AN patients and AN model animals), AgRP neuronal suppression triggered by movement would be an innate driver of exercise in AN patient. While effective treatments for the symptoms of AN have not yet been established^51–53^, selective control of AgRP neuronal activity may be developed to address critical symptoms of AN.

Walking has significant beneficial effects on health impacting the cardiovascular system, systemic metabolism (type diabetes), cancer and many other ailments^54^. A recent paper analyzed the impact of daily step count in the human population in relation to health benefits^54^. In this regard, our studies raise the possibility that the beneficial effects of walking on health are mediated, at least in part, by the hypothalamic melanocortin system. Future studies will be needed to confirm that observations we found in a murine model corresponds to the human condition, and whether manipulation of the melanocortin system influences the impact of walking on peripheral tissues, such as the liver, pancreas and heart.

## Methods

### Subjects

For this study, 4 female and 7 male AgRP-IRES-Cre mice (strain 012899, Jackson Laboratory; aged 9-16 weeks) and 2 female POMC-Cre mice (strain 005965, Jackson Laboratory; aged 33-42) were used (see Table 1). Mice were sourced from the Jackson Laboratory and maintained on a C57BL/6 genetic background. All experimental procedures were approved by the Institutional Animal Care and Use Committee (IACUC) of Yale University. The animals were housed in temperature- (20–23 °C) and humidity-(30-70%) controlled rooms, on a 12/12-hour light/dark cycle, with lights on from 07:00 to 19:00. Prior to the behavioral and fiber photometry experiments, the mice were moved to a reversed light cycle room, with lights on from 21:00 to 09:00, at least one week before the start of the experiments. Mice were provided with food and water ad libitum, except on specific days when behavioral experiments were conducted (as described below).

**Table 1.** Mice list used for *in-vivo* fiber photometry.

| ID | Genotype | Sex | Week-Old @ Surgery | Week-Old @ Exp | Wheel-Walking | Pavlovian | Feeding |
| --- | --- | --- | --- | --- | --- | --- | --- |
| A | POMC-Cre | Female | 33 | 42 | Yes | No | No |
| B | POMC-Cre | Female | 33 | 42 | Yes | No | No |
| C | POMC-Cre | Female | 33 | 42 | Yes | No | No |
| D | POMC-Cre | Female | 29 | 32 | Yes | No | No |
| E | POMC-Cre | Female | 29 | 32 | Yes | No | No |
| F | POMC-Cre | Female | 29 | 32 | Yes | No | No |
| G | AgRP-Cre | Male | 12 | 14 | Yes | Yes | Yes |
| H | AgRP-Cre | Male | 12 | 14 | Yes | Yes | Yes |
| I | AgRP-Cre | Male | 10 | 12 | Yes | Yes | Yes |
| J | AgRP-Cre | Male | 15 | 17 | Yes | Yes | Yes |
| K | AgRP-Cre | Female | 9 | 11 | Yes | Yes | Yes |
| L | AgRP-Cre | Female | 9 | 11 | Yes | Yes | Yes |
| M | AgRP-Cre | Male | 10 | 12 | Yes | Yes | Yes |
| N | AgRP-Cre | Male | 10 | 12 | Yes | Yes | Yes |
| O | AgRP-Cre | Female | 12 | 14 | Yes | Yes | Video file was broken |
| P | AgRP-Cre | Male | 16 | 18 | No | No | Yes |
| Q | AgRP-Cre | Female | 16 | 18 | No | No | Yes |

**Table 2.** Mice list used for *in-vivo* optogenetics.

| ID | Genotype | Opsin | Sex | Week-Old @ Surgery | Week-Old @ Exp | Optogenetics in Stay | Optogenetics in Walk |
| --- | --- | --- | --- | --- | --- | --- | --- |
| A | AgRP-Cre | EGFP | Male | 26 | 35 | Yes | Yes |
| B | AgRP-Cre | Jaws | Male | 26 | 35 | Yes | Yes |
| C | AgRP-Cre | Jaws | Male | 25 | 34 | Yes | Yes |
| D | AgRP-Cre | EGFP | Male | 25 | 27 | Yes | Yes |
| E | AgRP-Cre | Jaws | Male | 25 | 27 | Yes | Yes |
| F | AgRP-Cre | EGFP | Male | 24 | 26 | Yes | Yes |
| G | AgRP-Cre | Jaws | Male | 24 | 26 | Yes | Yes |
| H | AgRP-Cre | EGFP | Male | 24 | 28 | Yes | Yes |
| I | AgRP-Cre | Jaws | Male | 24 | 28 | Yes | Yes |
| J | AgRP-Cre | EGFP | Male | 18 | 22 | Yes | Yes |
| K | AgRP-Cre | Jaws | Male | 18 | 22 | No | Yes |
| L | AgRP-Cre | EGFP | Male | 10 | 14 | No | Yes |
| M | AgRP-Cre | Jaws | Male | 10 | 14 | No | Yes |
| N | AgRP-Cre | EGFP | Male | 10 | 14 | No | Yes |
| O | AgRP-Cre | EGFP | Male | 14 | 16 | No | Yes |
| P | AgRP-Cre | Jaws | Male | 14 | 16 | No | Yes |
| Q | AgRP-Cre | Jaws | Male | 20 | 22 | No | Yes |
| R | AgRP-Cre | Jaws | Male | 18 | 20 | No | Yes |

### Surgical procedures

All stereotaxic surgeries were performed under isoflurane anesthesia (4% for initial sedation, maintained at 1-2%) with regular monitoring to ensure a stable respiratory rate and absence of tail or toe pinch response. Mice were positioned in a stereotactic apparatus, and a heating pad was used to prevent hypothermia. Regional anesthesia was administered with an injection of 80 µL/10 g body weight of 0.5% Bupivacaine (Hospra), followed by 50 µL of Ethiqa XR (Fidelis Animal Health) for analgesia. After shaving the skull hair, a midline incision was made, and the skull surface was cleaned of all connective tissue. Craniotomies were performed using a drill over the designated injection sites and/or optic fiber placement locations, guided by coordinates from the Mouse Brain Atlas.

A virus expressing calcium indicators (AAV9-syn-FLEX-jGCaMP8m-WPRE: 162378-AAV9, Addgene) or mixture with viruses expressing inhibitory opsin (AAV5-CAG-FLEX-rc [Jaws-KGC-GFP-ER2]: 84445-AAV5, Addgene) or control fluorescent (AAV5-CAG-FLEX-EGFP-WPRE: 51502-AAV5, Addgene) was injected into arcuate nucleus (a single hemisphere for fiber photometry only or both hemispheres for optogenetics) at the following coordinates relative to the bregma: AP, -1.58 mm; ML, ±0.1 mm; DV, 5.7 mm. The injections were conducted using a nano injector (Nanoject III, Drummond) equipped with glass pipettes, creating a 50 µm pocket below the dorsoventral coordinate. After injection, the glass pipette was left in place for at least 5 minutes before retraction to allow for proper diffusion. A total of 900 nL of the virus solution was injected at a rate of 1 nL per second. For fiber photometry recordings, optic fibers (400 µm in diameter, 0.48 NA, Neurophotometrics) were implanted at the coordinates AP, -1.58 mm; ML, ±0.25 mm; DV, 5.65 mm, positioning the fiber tip 50 µm above the virus injection site. For optogenetics (with fiber photometry), the optoc fibers were implanted at AP, -1.58 mm; ML, ±0.25 mm; DV, 5.50 mm. The fibers were secured to the skull using a stereotaxic cannula holder (XCL, Thorlabs), along with Optibond (Kerr Dental) and Charisma (Heraeus Kulzer) dental materials. For head-fixation, a metal bracket was implanted on the skull’s surface using Charisma, and a final layer of dental cement was applied to secure the fibers and brackets.

Following surgery, mice were administered 100 µL/10 g body weight of Meloxidyl (Ceva Animal Health) as an antibiotic. Additional doses of Metacam were given 24 and 48 hours post-surgery. Mice were allowed to recover for at least 2 weeks before starting the experiments to ensure full recovery and adequate viral expression.

### Behavioral experiment

A total of 9 mice (3 females and 6 males) were used in the following sequence (Table S1): head-fixed wheel walking, head-fixed Pavlovian conditioning tasks, and free-moving feeding tests. Data from one mouse in the free-moving feeding test were excluded from the analysis due to a corrupted video file. An additional 2 mice (1 female and 1 male) were used exclusively for the free-moving feeding test (Table S1). The free-moving feeding test was conducted between 12:00 and 15:00, while the head-fixed wheel walking and Pavlovian conditioning tasks were performed between 09:00 and 12:00 under a dark cycle.

#### Fasting and water restriction

To ensure the activation of AgRP neurons during behavioral experiments involving fiber photometry and optogenetic manipulation, all food pellets were removed from the mice’s home cages in the afternoon (14:00-16:00) one day prior to the experiment (including the free-moving feeding test, head-fixed wheel walking after 3-day acclimation, test sessions of the Pavlovian conditioning task, and optogenetics). During the fasting period, water was available ad libitum in the home cage.

For the Pavlovian conditioning task training, daily water intake was mildly restricted. Mice received a maximum of 0.75 mL of water during each training session. Additionally, 1-2 mL of water was provided in their home cages daily to maintain their body weight above 85% of the original weight. During the period of water restriction, food was available ad libitum in the home cage.

#### Free-moving feeding test

The apparatus consisted of a Plexiglass open field (35 cm x 35 cm) and two types of wire cups (sensing and feeding cups). A USB camera was positioned above the open field to record the mice’s behavior (Extended Data Fig.1A). The experimental session comprised three blocks. Mice were first placed in a corner (starting point) of the open field for 10 minutes while remaining in their home cage (“control block”). Afterward, the mice were returned to their home cage for 2-3 minutes. A sensing cup (a metal-wire cup containing food pellets that the mice could not access; 5 cm in radius and 20 cm high; Extended Data Fig.1B left) was then placed in the diagonal corner opposite the starting point.

The mice were again placed at the starting point and observed in the open field for 10 minutes (“sensing block”). Following the sensing block, the mice were returned to their home cage for another 2-3 minutes. The sensing cup was then replaced with a feeding cup (a similar metal-wire cup containing food pellets that the mice could access; Extended Data Fig.1B right), and the mice were returned to the open field for an additional 10 minutes (“feeding block”). The experiment was conducted under dim lighting during the dark cycle and was performed with the mice in a fasted state. ***Head-fixed wheel walking***

The head-fixed wheel walking apparatus was based on the design described by Heiney et al.^21,22^. The wheel consists of a foam cylinder mounted on bearings that hold a metal axle, which is secured to a metal frame. The head bracket is attached to the frame using two screws, keeping the mouse’s head fixed on top of the cylinder while allowing the mouse to walk freely (Fig. 1A). Walking activity was recorded using a rotary encoder, with the analog signal digitized at 4 kHz via a USB-6001 data acquisition board (National Instruments) and recorded using a customized program in Bonsai software (https://www.open-ephys.org/bonsai). The rotary encoder signal was calibrated to convert voltage amplitude into speed (meters per second). The head-fixed wheel was placed inside a sound attenuation box to minimize external noise.

During the experimental session, mice were acclimated to the head-fixed wheel over three days, with sessions lasting 10, 20, and 30 minutes, respectively. One day before the experimental session, food pellets were removed from the mice’s home cages, and the mice were kept in a fasted state. For the wheel walking experiment with fiber photometry, mice were habituated on the head-fixed wheel for 2 minutes to prevent a significant reduction in fiber photometry background signal during the initial minutes. During this habituation phase, the wheel axis was locked to prevent movement. After the habituation phase, the wheel axis was unlocked, and BONSAI recording was conducted for 30 minutes.

#### Head-fixed wheel walking with optogenetics manipulation

In experiments involving optogenetic manipulation during head-fixed physical activity (i.e., immobility or walking), the same wheel apparatus described in the previous “head-fixed wheel walking” section was used. However, opto-laser control and data acquisition were managed using custom Python-based programs (https://www.python.org/) interfaced with a National Instruments data acquisition board (NI-DAQ6001).

To acclimate mice to the apparatus, they were first habituated to head-fixation over two days. On the first day, mice underwent a 20-minute session consisting of 10 minutes with the wheel locked, followed by 10 minutes of free walking. On the second day, mice experienced a 30-minute session consisting of 10 minutes with the wheel locked followed by 20 minutes of free walking. Food was removed from the home cage one day prior to the experiment to induce a fasted state.

For the optogenetic mimic of AgRP neuronal suppression with fiber photometry, mice were first habituated for 10 minutes on the head-fixed, locked-wheel apparatus prior to recording, to minimize baseline signal shifts in fiber photometry. Following habituation, the wheel was unlocked. During the first 10 minutes of the experimental session, only wheel movement was measured and recorded. In the subsequent 20 minutes, wheel movement was recorded as before, and opto-laser stimulation was triggered if the mouse remained immobile for more than 2 seconds. Each laser stimulation consisted of 4 seconds of irradiation followed by an inter-stimulation interval that varied randomly between 20 and 30 seconds.

For the optogenetic amplification of movement-driven AgRP neuronal suppression, mice were also habituated for 10 minutes on the locked wheel prior to the experimental session. After the session began, the wheel was unlocked to allow spontaneous walking. When walking was detected (defined as wheel velocity exceeding 0.02 m/sec), laser stimulation was administered in a pattern of 4 seconds of irradiation followed by a 1-second interval. This stimulation continued as long as the mouse remained walking, throughout the 30-minute session.

This optogenetic stimulation protocol was repeated for three consecutive days. To evaluate post-manipulation effects, a session without laser stimulation was conducted on the day following the final optogenetic session.

#### Head-fixed Pavlovian conditioning task

For the head-fixed Pavlovian conditioning task, we utilized the head-fixed wheel apparatus with the axis locked, serving as a stage for the mice during Pavlovian conditioning. The mice’s heads were fixed in the same manner as described for the head-fixed wheel experiment. A spout was positioned in front of their mouths (Fig. 2A) to deliver a 10% sucrose solution as a liquid food reward in response to the mice’s behavior. This spout also functioned as a lickometer, detecting licking based on fluctuations in resistance and providing feedback to a task control device during sessions. The task control device was built using Arduino, and its control software was created with a customized Python program (https://www.python.org/). A high-speed camera, recording at 100 Hz, was placed close to the mice’s faces to capture licking behavior, which was later analyzed using a customized DeepLabCut program^29^. The licking data obtained were used in subsequent analyses. Behavioral performance and task log data were recorded using the customized Python and BONSAI programs.

The head-fixed Pavlovian conditioning task consisted of 10 sessions—test sessions on days 1, 5, and 10, and training sessions on days 2, 3, 4, 6, 7, 8, and 9. In a test session (Fig. 2A), each trial began when the mice refrained from licking for 2-3 seconds, with the exact duration randomized for each trial. At the start of each trial, a reward-predictive cue (a 4 kHz pure tone lasting 0.3 seconds) was presented. A spout delivering 5 µL of a 10% sucrose solution as a liquid food reward was positioned near the mouse’s mouth, with the reward delivered 1 second after the cue in 70% of the trials (reward trials). In the remaining 30% of trials, no reward was delivered after the cue (omission trials). Each trial ended and transitioned to the next after a 3-second inter-trial interval if the mice licked the spout within 1 second after the cue onset, ensuring no liquid-food residue remained on the spout when a new trial began. In the training sessions, each trial proceeded in the same manner as in the test sessions, but the session consisted exclusively of reward trials. A session continued for 30 minutes or until the mice completed 150 trials. Test sessions were conducted under fasted conditions, while training sessions were conducted under water-restricted conditions.

### Fiber photometry

Calcium indicator (i.e., GCaMP8m) signals from AgRP neurons were measured using a commercial fiber photometry system, the LUX RZ10X processor (Tucker-Davis Technologies). Recording and data acquisition were controlled by Synapse software (Tucker-Davis Technologies). To correct for motion artifacts in the recorded calcium indicator signals, two excitation wavelengths were utilized with 30 µW power intensity. The 465 nm LED, modulated sinusoidally at 534 Hz, excited calcium-dependent fluorescence (calcium indicator signal). Additionally, the 405 nm LED, the isosbestic wavelength for GCaMP, was modulated sinusoidally at 211 Hz to acquire calcium-independent fluorescence, serving as a control signal. Both wavelengths were combined through an optical fiber patch (Doric Systems) using a minicube (FMC4, Doric Systems) and focused onto a femtowatt photoreceiver. The recorded signal was corrected using the control signal and synchronized with the behavioral experiment data (details are provided in the “Data Analysis” section). Data were acquired at a rate of 1 kHz.

### Optogenetics

To conduct laser irradiation according to walking behavior (i.e., wheel movement), opto-laser system was built as a closed-loop system that was controlled by a custom Python program monitoring wheel movement via the NI-DAQ6001 (Fig. 5a). 630 nm wavelength with 3 mW laser intensity was used for optogenetics stimulation. The custom Python program monitored wheel movements every moment and judged whether the mice were immobile or walk. This judgement was utilized for trigger timing of opto-laser (*Judgement criteria was dependent on section “Head-fixed wheel walking with optogenetics manipulation” for details).

### Data analysis

Data analysis for fiber photometry and behavioral experiments was conducted using customized Python programs, unless otherwise noted.

#### Fiber photometry

The recorded calcium indicator signal (coming from 465 nm LED excitation) was corrected using the control signal (coming from 405 nm LED excitation) by a modified protocol described previously in Martianova et al.^55^. To remove the slope and low-frequency fluctuations in signals, fit curves were calculated and subtracted from both calcium indicator and control signals separately (Extended Data Fig.4A). The signals were standardized using the mean value and standard deviation (z-scoring: z-scored signal = ( signal - average ) / SD; zSig465 and zSig405) (Extended Data Fig.4B). Using non-negative robust linear regression, fit standardized signals (Extended Data Fig.4B) were fit to the regression function: zSig465 = a * zSig405 + b (Extended Data Fig.4C). New values of zSig405 fitted to zSig465 (fitSig405) were found using the parameters a and b of the linear regression (fitSig405 = a * zSig405 + b) (Extended Data Fig.4D). The normalized dF/F (z-dF/F) was calculated (z-dF/F = zSig465 - fitSig405) (Extended Data Fig.4E). This normalized dF/F was additionally corrected at each behavioral event to show signal dynamics more precisely as described in the following analyses.

#### Free-moving feeding test

Objects, including the mice and sensing/feeding cups, were tracked throughout the analysis using a customized DeepLabCut program. Each block initiation was defined as follows: for the control block, it was the first entry of a mouse into the open field; for the sensing and feeding blocks, it was the first entry of a mouse into the open field with each respective cup present.

The baseline period for analysis was defined as the time window from -15 to -5 seconds relative to each block initiation. Standardized (z-scored) signals were calculated for each block within this baseline period. The time window from -5 to 0 seconds was excluded from the baseline period due to potential noise introduced by transferring the mice from their home cage to the open field during this interval.

The magnitude of the fiber photometry signal was quantified as the mean signal within the time window from 0 to +300 seconds relative to block initiation.

#### Head-fixed wheel walking

Although the recording platforms differed between the fiber photometry-only sessions and those involving optogenetic manipulation with fiber photometry, the same data analysis pipeline was applied to both datasets as described below.

Analog signals from the rotary encoder and fiber photometry were down sampled (from 4000 Hz to 20 Hz and from 1000 Hz to 20 Hz, respectively). The rotary encoder analog signal was converted into velocity (m/sec). Walking initiation was defined as when the wheel velocity exceeded 0.02 m/sec in the forward direction for the first time within the last 3 seconds, remained consecutively above this threshold for more than 3 seconds. Walking termination was defined as when the wheel velocity fell below 0.02 m/sec following walking initiation and it lasted for more than 3 seconds.

The fiber photometry signal was standardized (z-scored) using the mean and standard deviation from the baseline period between -2 to 0 seconds relative to walking initiation. This baseline period was also used for standardization at walking termination. Standardization was applied to each walking event.

For each walking initiation/termination, the average velocity and the fiber photometry signal within each time window (walking initiation pre-window, -2 ∼ 0 seconds from the initiation; walking initiation post-window, 0 ∼ 3 seconds; walking termination pre-window, -3 ∼ 0 seconds from walking termination; walking termination post-window, 0 ∼ 3 seconds from walking termination) were calculated. The median of these values across all walking events was used as the representative value for each experimental session.

#### Head-fixed Pavlovian conditioning task (Test session)

Licking responses were identified using a customized DeepLabCut program, with the timing of the tongue movement defined as the timing of the licking response. Both licking data and the measured fiber photometry signals were downsampled to 20 Hz.

The fiber photometry signal was standardized (z-scored) using the mean and standard deviation from a baseline period between -2 to 0 seconds relative to trial initiation. This baseline period was also used for standardizing the signal at reward detection. Standardization was performed for each trial.

For quantification during the cue window (between -2 to 0 seconds from cue onset), the licking number and the average of the fiber photometry signal were calculated as measures of licking and signal magnitude. For quantification during the reward detection window (between 0 to +1.5 seconds from the first licking response after reward delivery), the minimum fiber photometry signal within the window was used as the amplitude. The licking number was quantified for both the cue and reward detection windows. This quantification was performed on a trial-by-trial basis.

For estimation of the rate of change in AgRP neuron activity at the last licking, averaged AgRP neuron activity was calculated session-by-session at first. The difference was calculated using the averaged activity every time step. The difference was grouped into pre-/post-window (pre-window is between -2 to 0 seconds from the last lick; post-window is between 0 to 2 seconds from the last lick). Data were pooled according to trial type (i.e., reward or omission; fast or slow response; post-reward or post-omission) for each session. The median of the pooled data for each session was used as the representative value for the experiment.

#### Comparison among different behavioral tests

To compare AgRP neuron activity across head-fixed wheel walking, head-fixed Pavlovian conditioning (reward trials only), and free-moving feeding tests (feeding block only), 8 of 11 mice were included in the analysis. Three mice were excluded: one due to a lack of free-moving feeding test data caused by video recording issues, and two because they were not used for head-fixed behavioral tests (see Table S1). To quantify the magnitude of AgRP neuronal suppression, the average and minimum size of AgRP neuron activity were calculated within specific time windows. For head-fixed wheel walking and Pavlovian conditioning tasks, the time window from 0 to 4 seconds after walking initiation or cue onset was used. For the free-moving feeding test, the time window from 0 to 300 seconds after block initiation was analyzed.

#### Statistical analysis

Non-parametric statistical tests (i.e., Wilcoxon signed-rank test (paired comparison test), Friedman test (repeated measures ANOVA), and Spearman rank-order correlation) were employed, except for the following multiple linear regression (MLR), as some data did not follow a normal distribution. In analysis of the MLR, MLR models were estimated using ordinary least squares. F-test was used to test whether the estimated MLR model fits a dataset better than a model with no predictor variables and t-test was used to test whether each predictor variable significantly contributes to dependent variable. Kolmogorov–Smirnov test (KS-test) followed by FDR Benjamini-Hochberg correction was used for testing difference in the accumulated moved distance. FDR Benjamini-Hochberg correction was also applied when the Wilcoxon signed-rank test for walking velocity and KS-test and total moved-distance comparisons were repeated throughout the consecutive four-day sessions in optogenetics amplification of movement-driven AgRP neuronal suppression. All statistical analyses were conducted using built-in modules/functions in Python. Data were expressed as the median[Q1, Q3], and a p-value < 0.05 was considered statistically significant, and p < 0.1 was considered a tendency.

### Histology

For post hoc histological analysis, mice were anesthetized with isoflurane and transcardially perfused with 0.1 M phosphate-buffered saline (PB) followed by 4% paraformaldehyde. Brains were then dissected and fixed in 4% paraformaldehyde at 4°C for over 24 hours. After fixation, brains were sectioned into 100 µm thick slices using a vibratome.

Sections were washed three times for 5 minutes each in PB, then incubated in a blocking solution containing 5% normal donkey serum and 0.2% Triton X-100 in PB (PBT) for 30 minutes at room temperature. Following blocking, sections were incubated overnight at 4°C with chicken anti-GFP antibody (Abcam) at a dilution of 1:1000. The next day, sections were washed three times for 5 minutes each in PBT and then incubated for 3 hours at room temperature with goat anti-chicken Alexa Fluor 488 (Invitrogen) at a dilution of 1:500. Finally, sections were coverslipped with Vectashield containing DAPI (H-1200, VectaLabs) and imaged using a confocal microscope (LSM880 Airy Scan, Zeiss).

## Acknowledgements

We thank A. DeSimone of Yale School of Medicine Support Services for crafting head-fixed wheel apparatus and Y. Yasumoto of Dept. of Comparative Medicine in Yale School of Medicine for technical support in post-hoc histology and microscope imaging. Research reported in this publication was supported by R01DK126447(TLH) and R01AG067329(TLH) from the National Institute of Health, JP25H01746(SS) and JP25H02624(SS) from the Grants-in-Aid for Transformative Research Areas (A).

## Author contributions

Conceptualization: JY, TLH

Methodology: JY, NS, SS

Investigation: JY

Funding acquisition: TLH, SS

Project administration: JY

Supervision: IL, THL

Writing – original draft: JY, TLH

Writing – review & editing: JY, NS, SS, IL, TLH

## Competing interests

The authors declare no competing interests.

## Data and materials availability

It should be addressed to J.Y. / T.L.H.

---

**Extended Data 1.**
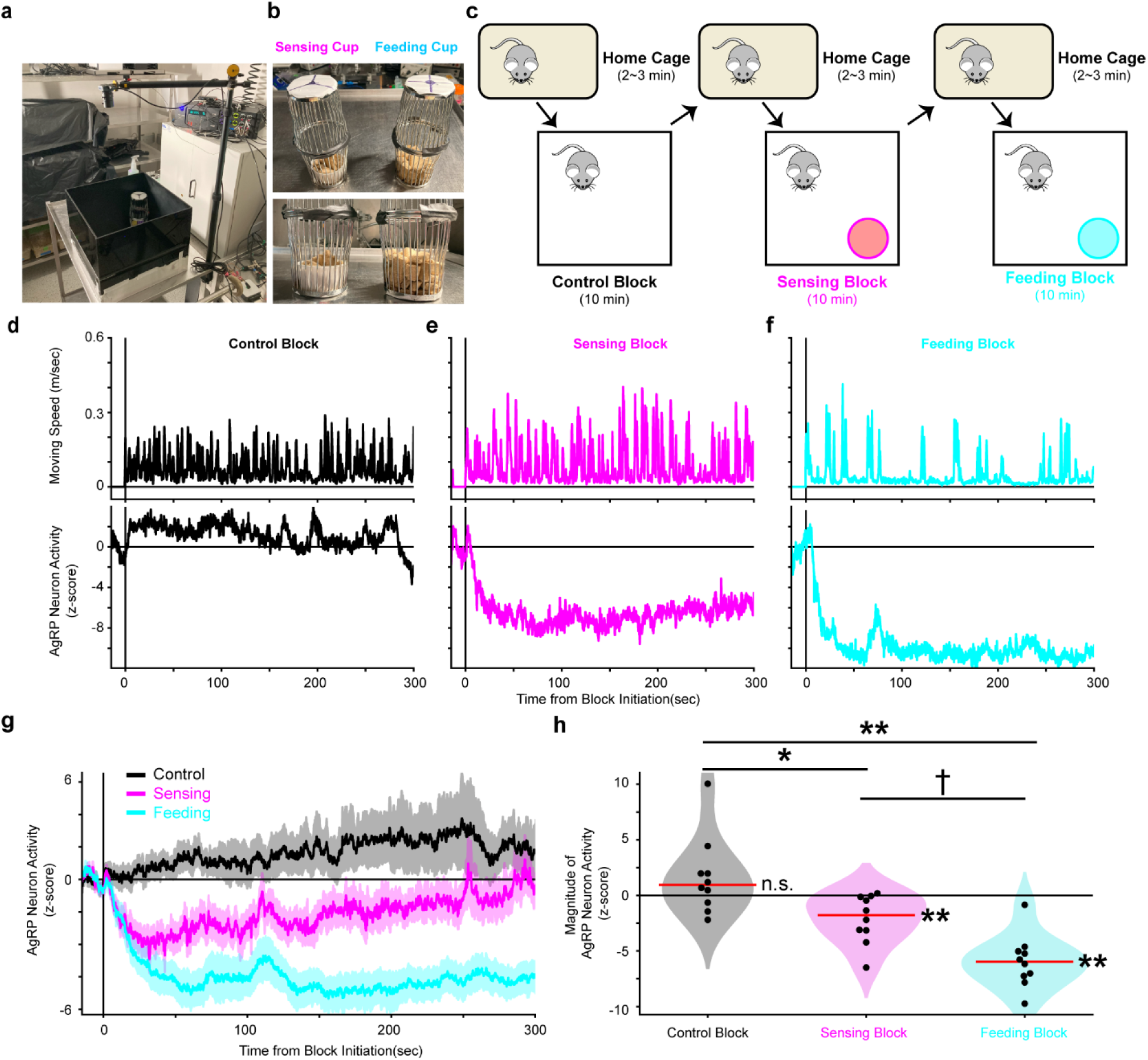
Food-sensing and feeding evoke continuous suppression in AgRP neurons. **(a)** Image of apparatus for free-moving feeding test. **(b)** Food sensing- and feeding cups. **(c)** Schematic of feeding test schedule. **(d∼f)** Example of moving velocity (top) and AgRP neuron activity (bottom) in control (d), sensing (e), and feeding blocks (f). **(g)** Average of AgRP neuron activity aligned at block initiation. **(h)** Distribution of AgRP neuron activity magnitude in each block. In plots, data is presented as mean ± SEM and bin time window is 100 ms. In violin plots, dots represent individual sessions and the red horizontal line indicates the median. ** next of the red line means p < 0.01 for Wilcoxon-signed rank test against 0. † between groups means p < 0.1, * between groups means p < 0.05 and ** between groups means p < 0.01 for post-hoc of the Friedman test to compare the groups. N = 10 mice.

**Extended Data 2.**
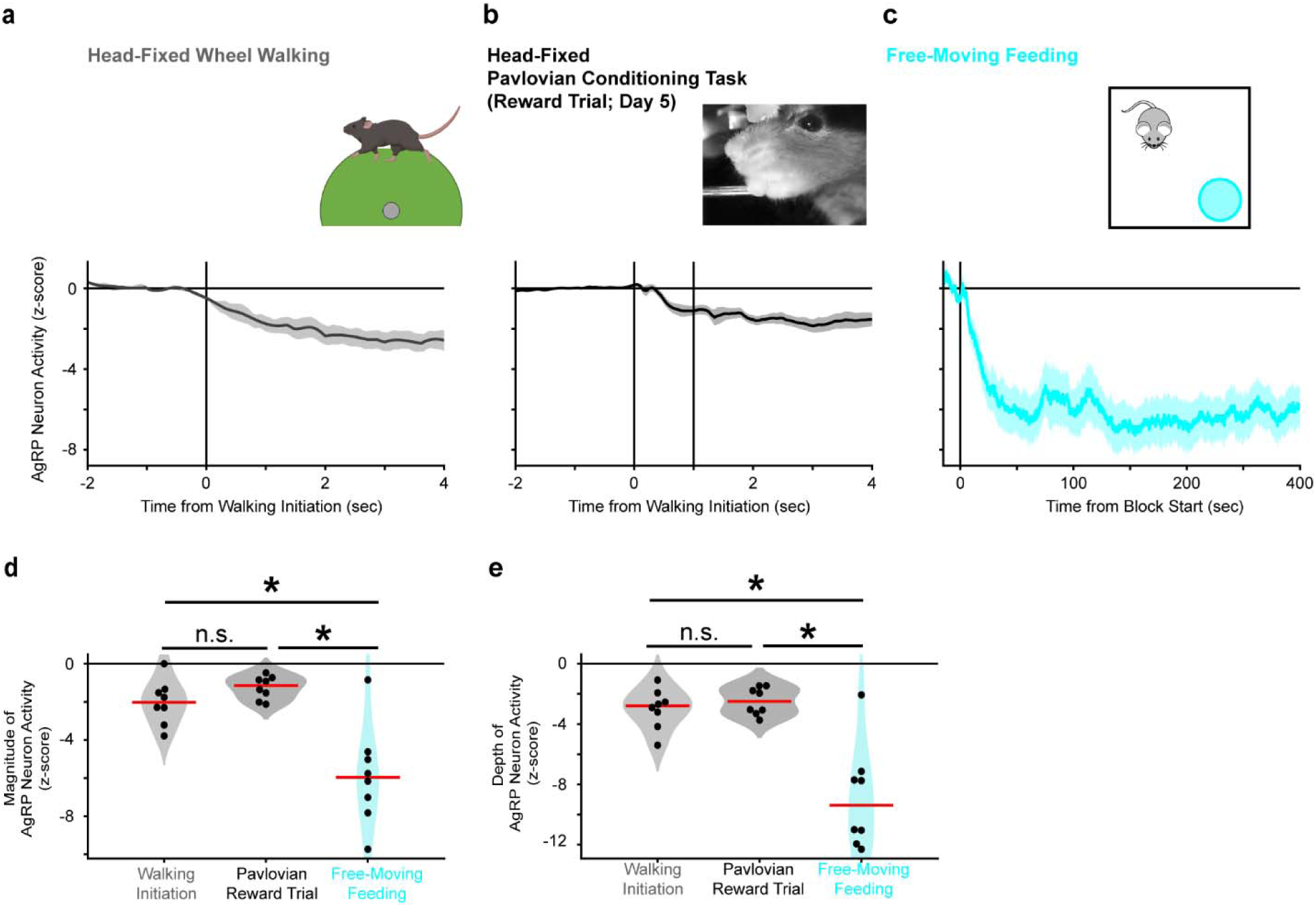
Movements trigger smaller suppression in AgRP neurons compared to feeding. **(a∼c)** Averaged AgRP neuronal suppression in head-fixed wheel walking (a), reward trial of head-fixed Pavlovian conditioning task (b), and feeding block during free-moving feeding test (c). **(d)** Distribution of mean size of the AgRP neuron activity. **(e)** Distribution of the minimum size of the AgRP neuron activity. In plots, data is presented as mean ± SEM. In plots of wheel walking and Pavlovian conditioning task, bin time window is 50 ms. And in plots of fee-moving feeding, bin time window is 100 ms. In violin plots, dots represent individual sessions and the red horizontal line indicates the median. * between groups means p < 0.05 for post-hoc of the Friedman test to compare the groups. N = 8 mice.

**Extended Data 3.**
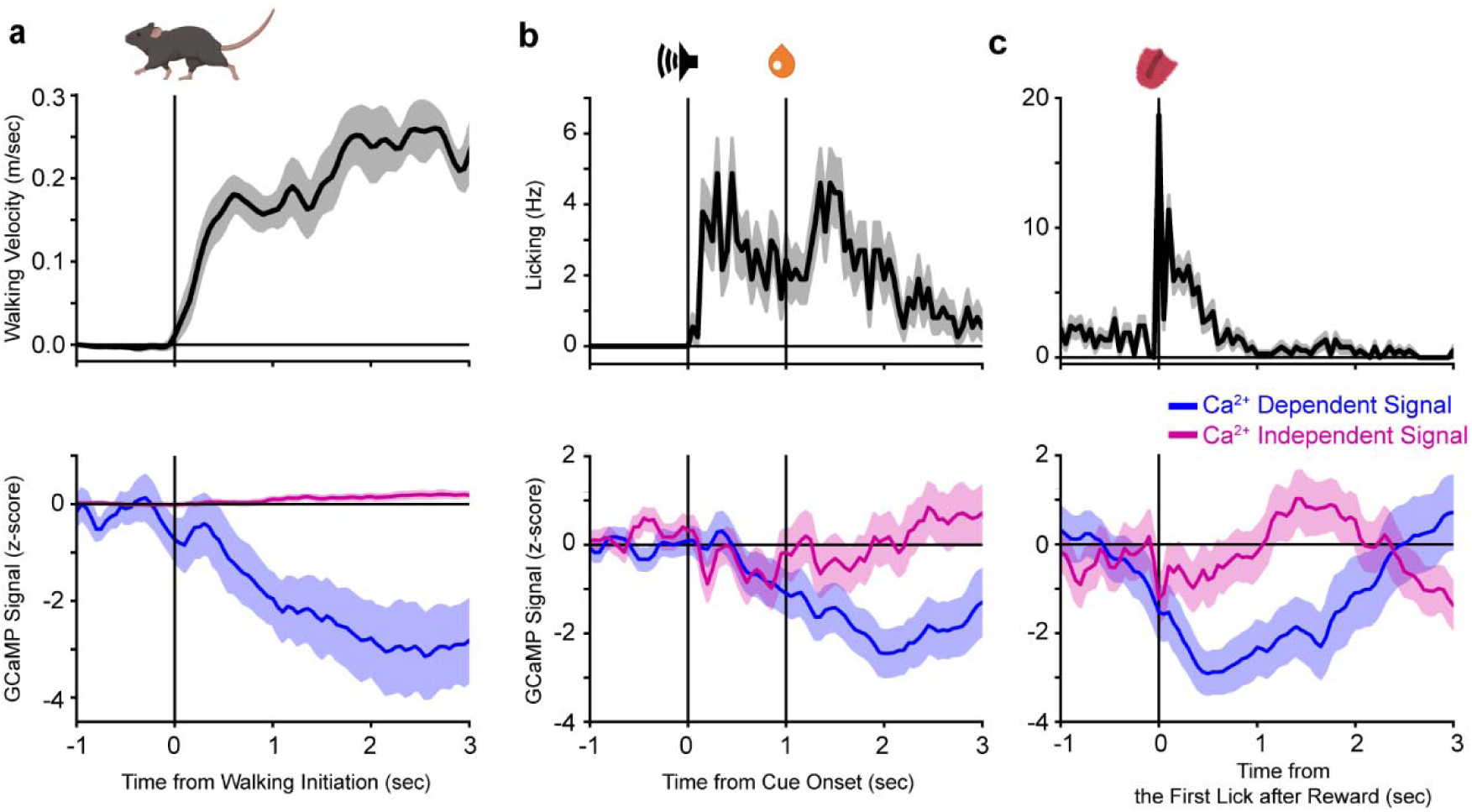
Movement-driven AgRP neuronal fluctuations are not motion artifact. **(a)** Average walking velocity (top) and Ca²^+^-dependent (blue) and Ca²^+^-independent (violet) signals (bottom) aligned to walking initiation in a representative mouse. **(b∼c)** Average licking frequency (top) and Ca²^+^-dependent (blue) and Ca²^+^-independent (violet) signals (bottom) aligned to cue onset (b) and the first lick after reward delivery (c) in a representative mouse. Data are presented as mean ± SEM with a bin size of 50 ms. N = 8 walking (a) and N = 74 reward trials (b).

**Extended Data 4.**
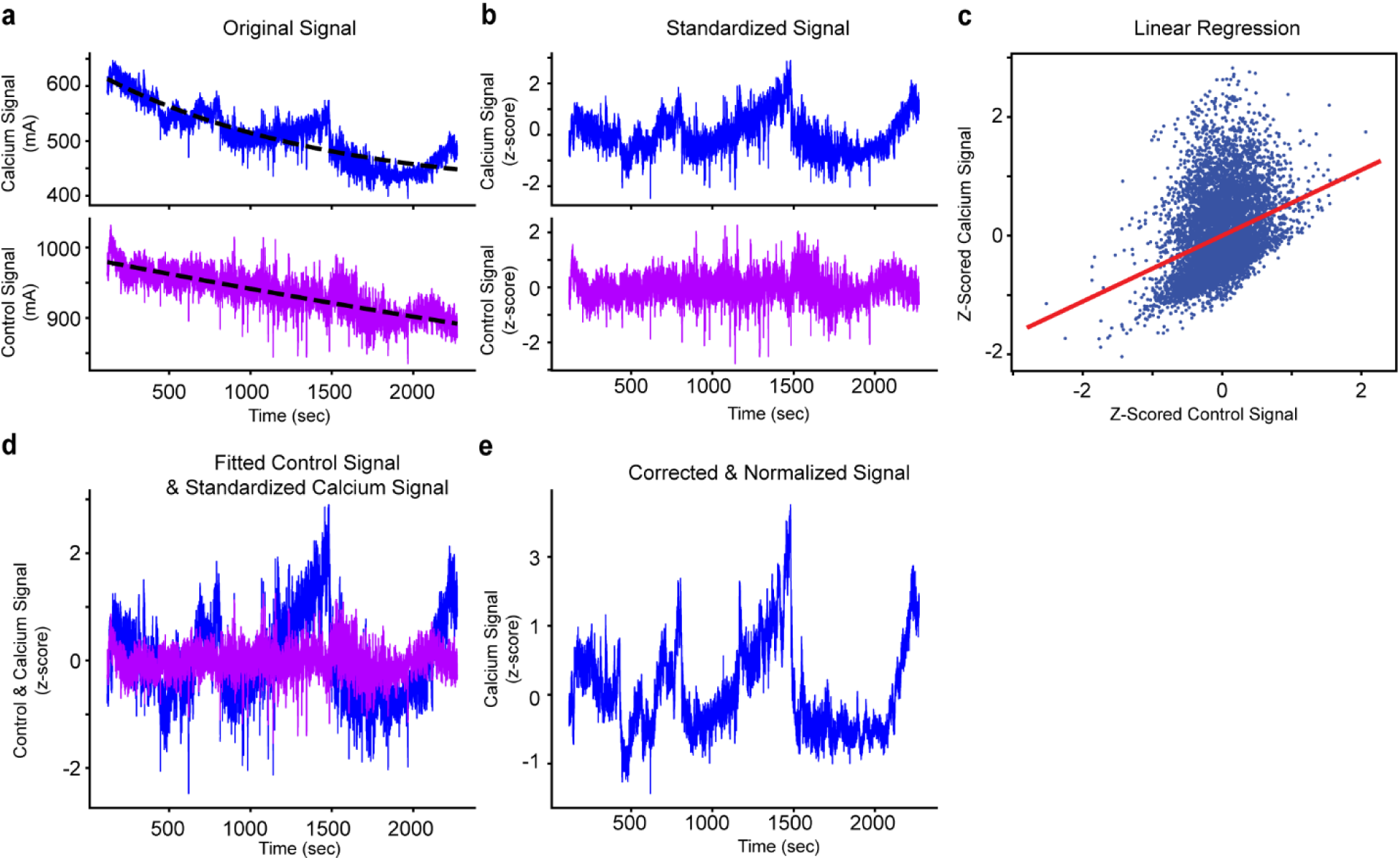
Standardization of fiber photometry signal using the isosbestic point wavelength. **(a)** Recorded original calcium indicator (excited by 465 nm LED) and control (excited by 405 nm LED) signals. Black lines indicate the baseline determined using a fit-curve algorithm. **(b)** Standardized signals after baseline correction. **(c)** Non-negative robust linear regression of calcium indicator and control signals. **(d)** Alignment of the standardized calcium indicator signal (zSig465) with the fitted control signal (fitSig405). **(e)** Corrected and normalized calcium-dependent signal.

**Extended Data 5.**
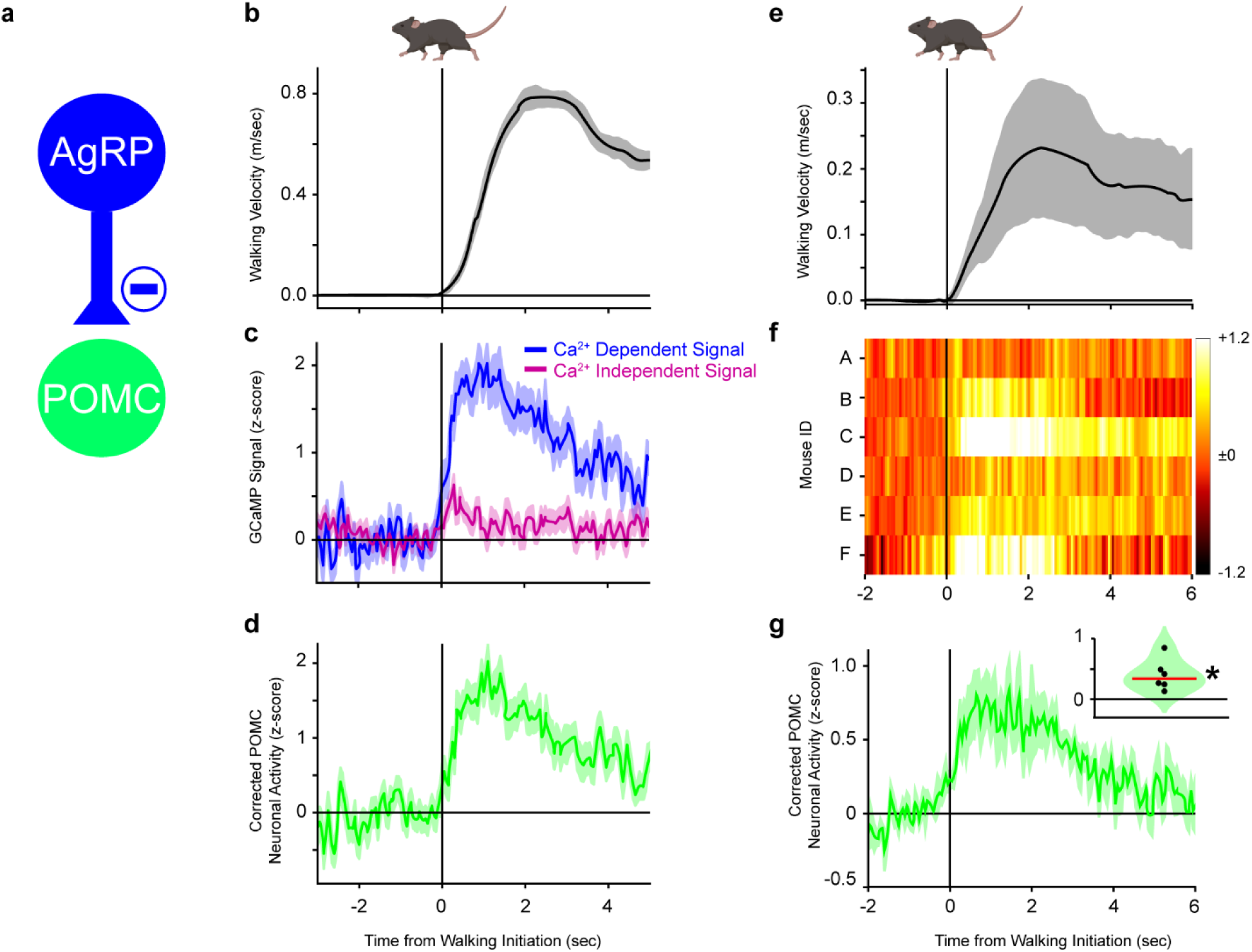
Walking initiates rapid activation of arcuate POMC neurons. (a) Schematic illustrating the known inhibitory projections from AgRP neurons to POMC neurons within the arcuate nucleus of the hypothalamus. (b–d) Data from a representative POMC-Cre mouse showing (b) average walking velocity, (c) Ca²^+^-dependent (blue) and Ca²^+^-independent (violet) fiber photometry signals, and (d) corrected POMC neuron activity (Ca²^+^-dependent signal normalized by the Ca²⁺-independent signal), all aligned to walking initiation. (e–g) Summary across animals showing (e) average walking velocity, (f) heat map of corrected POMC neuron activity in individual mice, and (g) average corrected POMC neuron activity aligned to walking initiation. The inset in (g) shows the distribution of POMC activity after walking onset. * means p < 0.05 for Wilcoxon-signed rank test against 0. Data are shown as mean ± SEM using 50 ms time bins. n = 39 walking bouts for panels b–d; N = 5 mice for panels e–g.

**Extended Data 6.**
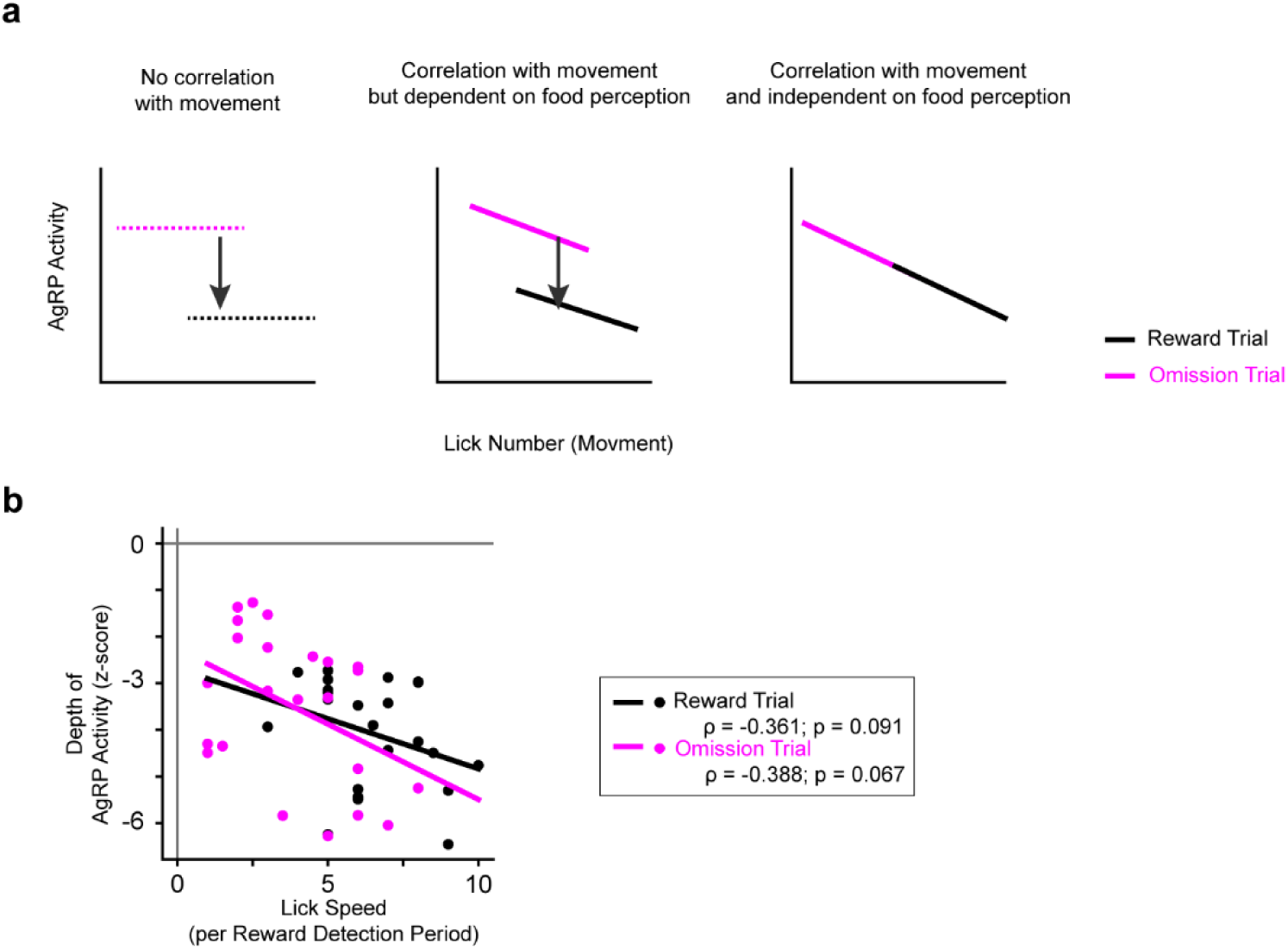
Licking-signal correlations in reward/omission trials are not significantly different. **(a)** Expected correlations between AgRP neuronal activity and licking number (movement) for each hypothesis. **(b)** Correlation between licking number and AgRP neuron activity amplitude. In the correlation plots, dots represent individual session and solid lines represent correlation line. The correlation lines, ρ, and p are estimated by Spearman rank-order correlation. N = 9 mice and n = 23 sessions.

**Extended Data 7.**
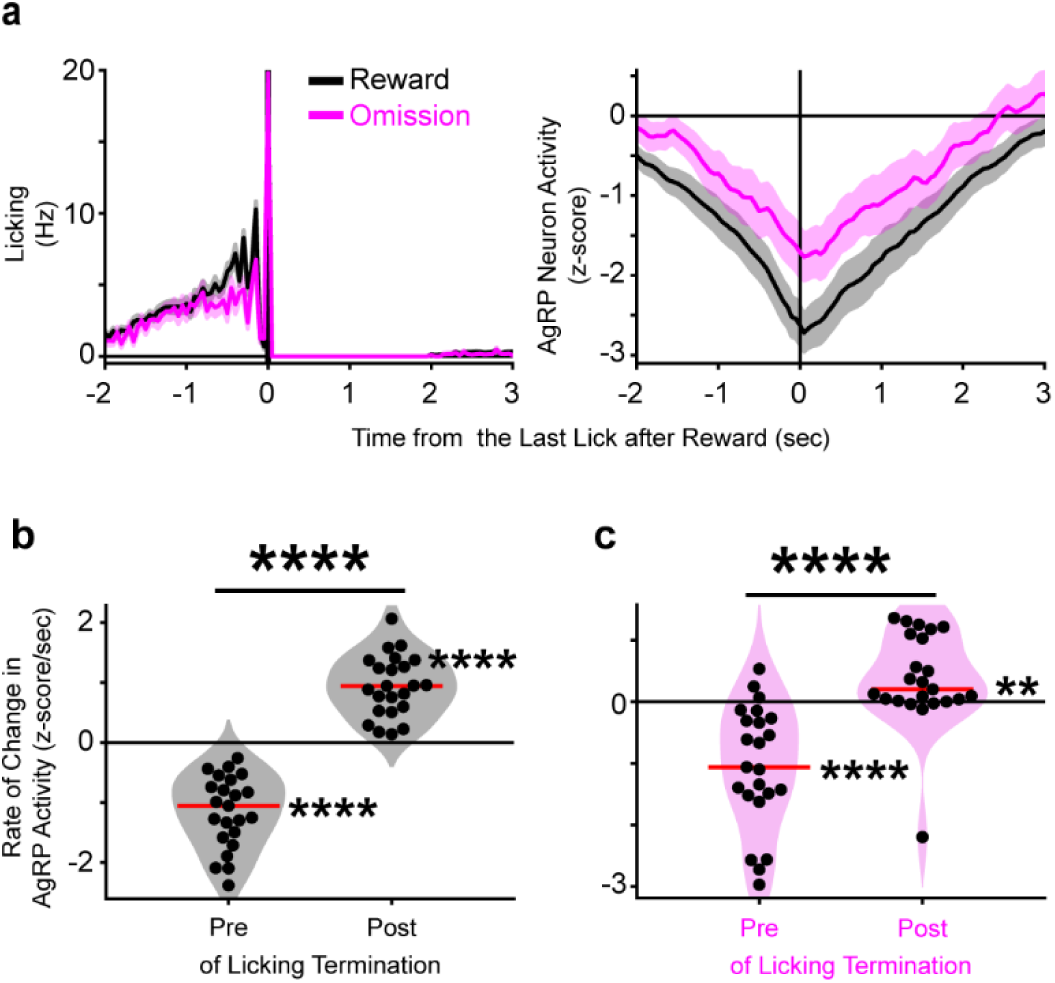
Licking-triggered AgRP neuronal suppression ends at the termination of the licking bout. **(a)** Average licking (left), and AgRP neuron activity (right) aligned at the last licking response after reward delivery. **(b∼c)** Distribution of the rate of change in AgRP neuron activity in pre-/post-window in reward trials (b) and omission trials (c). In plots, data is presented as mean ± SEM and bin time window is 50 ms. In violin plots, dots represent individual session, and the red horizontal line indicates the median. **** next of the red line means p < 0.0001 and ** means p < 0.01 for Wilcoxon-signed rank test against 0. **** between groups means p < 0.0001 for Wilcoxon-signed rank test to compare the pre- and post-windows. N = 9 mice and n = 23 sessions.

**Extended Data 8.**
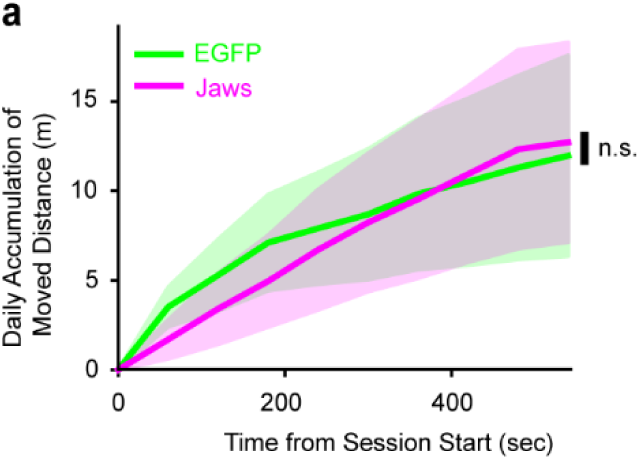
Walking performance was not significantly different between AgRP-Jaws and AgRP-EGFP mice before optogenetics manipulation. Accumulated walking distance on before optogenetics manipulation (Day -1). Green indicates AgRP-EGFP mice and magenta indicates AgRP-Jaws mice. Data is shown as mean ± SEM using a time bin. N = 8 AgRP-EGFP mice and 10 AgRP-Jaws mice.

## Notes

### Competing Interest Statement

The authors have declared no competing interest.

